# Exploring novel fluorine rich fuberidazole derivatives as hypoxic cancer inhibitors: design, synthesis, pharmacokinetics, molecular docking and DFT evaluations

**DOI:** 10.1101/2022.01.06.475235

**Authors:** Muhammad Babar Taj, Omar Makarm Ali

## Abstract

Sixteen fuberidazole derivatives as potential new anticancer bioreductive prodrugs are prepared and characterized. The in vitro anticancer potential is examined to explore their cytotoxic properties employing apoptosis, DNA damage, and proliferation tests on chosen hypoxic cancer cells. Overall, eight substances (Compound **5a, 5c, 5d, 5e, 5g, 5h, 5i**, and **5m**) showed good cytotoxic properties. The potential of compounds is also examined through in silico studies (against *human serum albumin*), including chem-informatics to understand the structure-activity relationship (SAR), pharmacochemical strength, and the mode of interactions responsible for their action. The DFT calculations revealed that only **5b** compound showed the lowest ΔET (2.29 eV) while **5i** showed relatively highest βtot (69.89 × 10-31 esu), highest αave (3.18 × 10-23 esu), and dipole moment (6.49 Debye). This study presents a novel class of fuberidazole derivatives with selectivity toward hypoxic cancer cells.

## Introduction

Hypoxia plays a vital role in cancer cell survival against cancer therapy. Cancer cells show differences in cellular biological activities and the level of reactive oxygen species (ROS) due to higher metabolic activity(1). A high level of ROS disrupts mitochondrial function. The hypoxia condition is referred to the less oxygen supply at the tissue level, and cancer cells show increased antioxidant ability to defend ROS stress(2). That is why hypoxia causes unequivocal resistance to cancer treatments and suppresses the regulation of DNA repair pathways(3). Therefore, pharmacologically targeting hypoxia or ROS defence system, for example, the ability to form ROS and interaction with DNA, can cause mitochondria toxicity, which could mediate the death of cancer cells(4).

Controlling the survival effect of tumor cells ensures the effectiveness of targeted therapies. Researchers are trying to investigate the use of specific substances that could minimize the survival effect with the help of a bio-reductive mechanism in hypoxia conditions(5). Exploiting this concept, a wide range of different prodrugs were expanded, and their activation for selective cytotoxins has experimented, and interestingly, aniline derivatives were found to be the first class of bioreductive prodrugs(6). Currently, many chemical compounds containing quinones, heterocyclic N-oxides (CB 1954, tirapazamine, AQ4N), and nitro group are being used as radical prodrugs in cancer therapy(7). The most promising attribute of these chemical components is their potential to cause DNA damage by generating cytotoxic agents(8). The benzimidazole derivatives have been particularly important as potential agents for damaging the DNA of cancer cells and have been intensively investigated to explore their anticancer properties(9). In fuberidazole, the benzimidazole nucleus is categorized as a potential pharmacophore in medicinal chemistry, and its extraction from natural sources, such as cyanocobalamin that has further extended its pharmaceutical applications(10). It is well reported that compounds containing O–H bonds are reported to act as antioxidants; in addition, the N-H bonded amines also work as an antioxidant. They have garnered considerable study focus because cyclic amines have always been the primary structure of a large number of currently used medications(11). Benzimidazoles nucleus contained in fuberidazoles is a very popular nitrogen-containing heterocycle. They have anticonvulsant(12), antifungal(13), antiviral(14), and anticancer(15) activities. The structural resemblance of fuberidazole to the naturally occurring nucleotides allows it to target the polymers in the living world leading to a vast impact in biological systems(16).

Numerous recent studies suggested that the presence of nitrogen and oxygen functionalities in the fuberidazole compound plays a significant role in cultivating pharmacokinetics and pharmacodynamics traits of anticancer drugs by improving polarity, lipophilicity or other physicochemical properties(17).

Furthermore, the presence of nitrogen and oxygen functionalities in the fuberidazole (C_11_H_8_N_2_O) molecule gives its special characteristics to exhibit extended π-electron delocalization, which could be useful in organic electronics, organic light-emitting diodes (OLEDs)(18), solar cells(19), and nonlinear optical (NLO) applications(20).

Moreover, computational studies of fuberidazole derivatives were conducted to determine their pharmacodynamics (chemo-informatics and Lipinski’s rule validation), binding affinities via molecular docking, initial structure-activity relationships (SARs), and density functional theory (DFT) calculations to determine their triplet energies, dipole moments, polarizabilities, and first-order hyperpolarizabilities to analyze their NLO properties. We also tuned the substituents on both the donor and acceptor units of fuberidazole compounds and studied the effect of inserting π-bridge between them to enhance their NLO response.

Because of these facts, we intended to synthesize some novel fuberidazole derivatives as potential targets for DNA ruin at hypoxia MCF-7 cells. Additionally, we also analyzed their cytotoxic activity with the help of apoptosis and computational parameters.

## Materials and methods

All chemicals were obtained commercially and utilized without purification. Methanol, ethanol, and DMSO were procured from Sigma Aldrich, purified, and dried by standard analytical procedures. The melting points were determined using Gallon Kamp’s melting point instrument. The FT-IR spectrophotometer from Perkin Elmer was used to collect the infrared spectra. The ^1^H NMR (300 MHz) and ^13^C NMR (75.43 MHz) spectra were collected on a Bruker AM-250 spectrometer utilizing CDCl_3_ and DMSO as internal standards. A UV-Visible spectrophotometer was used to obtain UV-Visible absorption spectra, model UV-1700. Elements were analyzed using a Leco CHNS-932 Elemental Analyzer (Leco Corporation, USA).

### General method for the compounds (5a-p) synthesis

In the first step, the halo substituted furfural (10 mmol) was added in a stirred solution of trifluorobenzene-1,2-diamine (10 mmol) in a more environmentally friendly solvent polyethylene glycol (PEG_400_). The reaction mix was refluxed at an elevated temperature of 80-85 °C. TLC was used to monitor the reaction progress. Following aqueous workup, the crude product was obtained and refined with a mobile phase of hexane: ethyl acetate to get the pure compound (**3a**-**d**). In the second step, the compound **3a**-**d** (10 mmol) was stirred with equimolar NaH in acetone for one hour. After one hour, the appropriate alkyl halide (10 mmol) was added to the mixture and agitated for a further five hours at room temperature. TLC has also been used for monitoring the reaction progress. After the reaction was completed, the organic extract was prepared by dissolving the reaction mixture in water and extracting it with dichloromethane (3×100 mL). The organic extract was dried in the presence of anhydrous sodium sulphate and then imaged by chromatography over silica gel using ethyl acetate: petroleum ether (1:1) as the eluting system to obtain pure fuberidazole (**5a**-**p**) as a pure product. Following fuberidazole, derivatives were obtained by using the above procedure.

#### 4,5,6-trifluoro-2-(4-fluorofuran-2-yl)-1H-benzo[d]imidazole (5a)

Yield: 81%; m.p.: 149-153 °C; FT- IR ν (cm^-1^): 3296, 3035, 2953, 1621, 1567, 1543, 1446, 1458, 1423, 1327, 1128; ^1^H-NMR (300 MHz, DMSO-d_6_); δ (ppm) 11.49 (s, 1H, NH), 7.5 (s, 1H, =CH-O), 7.3 (m, 1H, Ar-H), 6.27 (d, 1H, furan J=6Hz); ^13^C-NMR (75 MHz, DMSO-d_6_) δ (ppm) 155.4, 159.7, 143.7, 141.3, 140.7, 138.2, 115.9, 115.1, 109.3, 108.2, 103.5; Anal. Calcd. for C_11_H_4_F_4_N_2_O: C, 51.58; H, 1.57; N, 10.94; found: C, 51.26; H, 1.84; N, 11.20.

#### 4,5,6-trifluoro-2-(4-fluorofuran-2-yl)-1-methyl-1H-benzo[d]imidazole (5b)

Yield: 70%; m.p.: 150-151 °C; FT- IR ν (cm^-1^): 3263, 3125, 2959, 1618, 1589, 1522, 1491, 1458, 1402, 1339, 1167; ^1^H-NMR (300 MHz, DMSO-d_6_); δ (ppm) 7.40 (s, 1H, =CH-O), 7.20 (s, 1H, Ar-H), 6.81 (s, 1H, furan), 3.63 (s, 3H, methyl); ^13^C-NMR (75 MHz DMSO-d_6_) δ (ppm) 155.45, 149.58, 144.24, 140.97, 139.67, 115.12, 114.61, 109.70, 107.21, 103.43, 29.98; Anal. Calcd. for C_12_H_6_F_4_N_2_O: C, 53.35; H, 2.24; N, 10.37; found: C, 53.26; H, 2.48; N, 10.55.

#### 1-ethyl-4,5,6-trifluoro-2-(4-fluorofuran-2-yl)-1H-benzo[d]imidazole (5c)

Yield: 60%; m.p.: 153-155 °C; FT- IR ν (cm^-1^): 3305. 3289, 3002, 2933, 1636, 1534, 1517, 1477, 1467, 1440, 1345, 1122; ^1^H-NMR (300 MHz, DMSO-d_6_); δ (ppm) 7.49 (s, 1H, =CH-O), 7.30 (s, 1H, Ar-H), 6.79(s, 1H, furan), 3.61 (m, 2H, -CH_2_), 2.51(t, 3H, - CH_3_, J=18Hz); ^13^C-NMR (75 MHz DMSO-d_6_) δ (ppm) 155.1, 150.09, 143.69, 141.5, 140.1, 138.3, 115.3, 114.7, 109.2, 108.1, 103.6, 33.7, 16.1; Anal. Calcd. for C_13_H_8_F_4_N_2_O: C, 54.94; H, 2.84; N, 9.86; found: C, 54.77; H, 3.11; N, 9.99.

#### 4,5,6-trifluoro-2-(4-fluorofuran-2-yl)-1-isopropyl-1H-benzo[d]imidazole (5d)

Yield: 76%; m.p.: 149-152 °C; FT- IR ν (cm^-1^): 3301, 3028, 2966, 1635, 1545, 1521, 1441, 1406, 1341, 1103; ^1^H-NMR (300 MHz, DMSO-d_6_); δ (ppm) 7.46 (s, 1H, =CH-O), 7.23 (m, 1H, Ar-H), 6.67(s, 1H. furan) 3.75(m, 2H, -CH_2_), 2.03(t, 6H, -CH_3_); ^13^C-NMR (75 MHz DMSO-d_6_) δ (ppm) 155.21, 149.97, 143.5, 141.32, 140.01, 138.33, 115.4, 114.4, 109.4, 107.53, 103.5, 45.12, 25.11; Anal. Calcd. for C_14_H_10_F_4_N_2_O: C, 56.38; H, 3.38; N, 9.39; found: C, 56.19; H, 3.26; N, 9.09.

#### 2-(4-chlorofuran-2-yl)-4,5,6-trifluoro-1H-benzo[d]imidazole (5e)

Yield: 82%; m.p.: 156-157 °C; FT- IR ν (cm^-1^): 3285, 3024, 2922, 1609, 1581, 1523, 1420, 1418, 1341, 1113; ^1^H-NMR (300 MHz, DMSO-d_6_); δ (ppm) 11.50 (s, 1H, NH), 7.47 (s, 1H, =CH-O), 7.23-7.25 (d, 1H, Ar-H, J=4.8Hz), 6.57(s, 1H, furan); ^13^C-NMR (75 MHz DMSO-d_6_) δ (ppm) 155.3, 150.1, 144.49, 142.3, 140.1, 135.98, 115.32, 114.71, 108.77, 101.22, 102.41; Anal. Calcd. for C_11_H_4_ClF_3_N_2_O: C, 48.46; H, 1.48; N, 10.28; found: C, 48.76; H, 1.63; N, 10.57.

#### 2-(4-chlorofuran-2-yl)-4,5,6-trifluoro-1-methyl-1H-benzo[d]imidazole (5f)

Yield: 81%; m.p.: 157-160 °C; FT- IR ν (cm^-1^): 3301, 3092, 2954, 1609, 1578, 1551, 1482, 1419, 1403, 1341, 1132; ^1^H-NMR (300 MHz, DMSO-d_6_); δ (ppm) 7.45 (s, 1H, =CH-O), 7.28 (s, 1H, Ar-H), 6.81 (s, 1H, furan), 3.31 (s, 1H, methyl); ^13^C-NMR (75 MHz DMSO-d_6_) δ (ppm) 155.42, 150.10, 143.29, 142.22, 140.11, 138.53, 115.21, 114.43, 109.20, 105.11, 25.1; Anal. Calcd. for C_12_H_6_ClF_3_N_2_O: C, 50.28; H, 2.11; N, 9.77; found: C, 50.47; H, 1.92; N, 10.01.

#### 2-(4-chlorofuran-2-yl)-1-ethyl-4,5,6-trifluoro-1H-benzo[d]imidazole (5g)

Yield: 65%; m.p.: 151-154 °C; FT- IR ν (cm^-1^): 3341, 3063, 2960, 1617, 1554, 1531, 1464, 1421, 1410, 1333, 1150; ^1^H-NMR (300 MHz, DMSO-d_6_); δ (ppm) 7.46 (d, 1H, =CH-O), 7.27 (s, 1H, Ar-H), 6.72(s, 1H, furan) 3.77 (m, 2H, -CH_2_), 2.51(t, 3H, -CH_3_, J=18Hz); ^13^C-NMR (75 MHz DMSO-d_6_) δ (ppm) 155.21, 150.43, 143.45, 141.27, 138.43, 115.5, 114.7, 109.5, 105.95, 103.89, 33.6, 16.1; Anal. Calcd. for C_13_H_8_ClF_3_N_2_O: C, 51.93; H, 2.68; N, 9.32; found: C, 52.11; H, 2.80; N, 9.50.

#### 2-(4-chlorofuran-2-yl)-4,5,6-trifluoro-1-isopropyl-1H-benzo[d]imidazole (5h)

Yield: 66%; m.p.: 157-159 °C; FT- IR ν (cm^-1^): 3233, 3091, 2947, 1630, 1568, 1532, 1452, 1434, 1420, 1328, 1109; ^1^H-NMR (300 MHz, DMSO-d_6_); δ (ppm) 7.26 (s, 1H, =CH-O), 7.2 (s, 1H, Ar-H), 6.49(s, 1H. furan) 4.78(m, 1H, -CH_2_), 1.81(t, 6H, -CH_3_); ^13^C-NMR (75 MHz DMSO-d_6_) δ (ppm) 154.89, 150.11, 140.2, 139.91, 136.12, 115.2, 114.4, 109.5, 106.81, 103.2, 45.23, 24.72; Anal. Calcd. for C_14_H_10_ClF_3_N_2_O: C, 53.43; H, 3.20; N, 8.90; found: C, 53.70; H, 3.31; N, 8.93.

#### 2-(4-bromofuran-2-yl)-4,5,6-trifluoro-1H-benzo[d]imidazole (5i)

Yield: 69%; m.p.: 153-156 °C; FT- IR ν (cm^-1^): 3296, 3035, 2953, 1621, 1567, 1543, 1446, 1458, 1423, 1327, 1128; ^1^H-NMR (300 MHz, DMSO-d_6_); δ (ppm) 11.00 (s, 1H, NH), 7.71 (s, 1H, =CH-O), 7.28 (d, 1H, Ar-H J=4.8Hz), 6.57 (s, 1H. furan); ^13^C-NMR (75 MHz DMSO-d_6_) δ (ppm) 154.96, 150.1, 143.4, 138.23, 136.11, 115.5, 114.9, 109.3, 108.2, ; Anal. Calcd. for C_11_H_4_BrF_3_N_2_O: C, 41.67; H, 1.27; N, 8.84; found: C, 41.95; H, 1.39; N, 9.11.

#### 2-(4-bromofuran-2-yl)-4,5,6-trifluoro-1-methyl-1H-benzo[d]imidazole (5j)

Yield: 77%; m.p.: 159-163 °C; FT- IR ν (cm^-1^): 3302, 3056, 2934, 1657, 1599, 1538, 1452, 1429, 1401, 1318, 1130; ^1^H-NMR (300 MHz, DMSO-d_6_) δ (ppm) 7.25 (s, 1H, =CH-O), 7.06 (s, 1H, Ar-H), 6.55 (s, 1H. furan), 3.42 (s, 3H, -CH_3_); ^13^C-NMR (75 MHz DMSO-d_6_) δ (ppm) 154.13, 149.7, 143.8, 140.9, 139.5, 136.4, 114.8, 109.1, 106.55, 103.4, 29.8; Anal. Calcd. for C_12_H_6_BrF_3_N_2_O: C, 43.53; H, 1.83; N, 8.46; found: C, 43.70; H, 2.09; N, 8.18.

#### 2-(4-bromofuran-2-yl)-1-ethyl-4,5,6-trifluoro-1H-benzo[d]imidazole (5k)

Yield: 59%; m.p.: 163-164 °C; FT- IR ν (cm^-1^): 3229, 3061, 2962, 1634, 1570, 1521, 1461, 1444, 1431, 1330, 1129; ^1^H-NMR (300 MHz, DMSO-d_6_); δ (ppm) 7.7 (s, 1H, =CH-O), 7.13 (s, 1H, Ar-H), 6.80(s, 1H, furan) 3.40 (m, 2H, -CH_2_), 2.47(t, 3H, -CH_3_, J=11Hz). ^13^C-NMR (75 MHz DMSO-d_6_) δ (ppm) 154.0, 150.4, 141.2, 143.1, 138.0, 136.0, 100.2, 105.5, 108.1, 33.2, 16.4; Anal. Calcd. for C_13_H_8_BrF_3_N_2_O: C, 45.24; H, 2.34; N, 8.12; found: C, 45.55; H, 2.64; N, 7.89.

#### 2-(4-bromofuran-2-yl)-4,5,6-trifluoro-1-isopropyl-1H-benzo[d]imidazole (5l)

Yield: 68%; m.p.: 160-162 °C; FT- IR ν (cm^-1^): 3296, 3035, 2953, 1621, 1567, 1543, 1446, 1458, 1423, 1327, 1128; ^1^H-NMR (300 MHz, DMSO-d_6_); δ (ppm) 7.48 (s, 1H, =CH-O), 6.77 (s, 1H, Ar-H), 6.48(s, 1H. furan) 3.85(m, 2H, -CH_2_), 1.62(s, 3H, -CH_3_), 1.85(s, 3H, -CH_3_). ^13^C-NMR (75 MHz DMSO-d_6_) δ (ppm) 155.2, 149.6, 142.9, 141.2, 139.9, 138.2, 115.4, 114.2, 108.5, 108.0, 103.2, 44.8, 24.5; Anal. Calcd. For C_14_H_10_BrF_3_N_2_O: C, 46.82; H, 2.81; N, 7.80; found: C, 47.11; H, 3.03; N, 7.64.

#### 4,5,6-trifluoro-2-(4-iodofuran-2-yl)-1H-benzo[d]imidazole (5m)

Yield: 80%; m.p.: 167-168 °C; FT- IR ν (cm^-1^): 3290, 3039, 2958, 1627, 1566, 1547, 1443, 1459, 1420, 1325, 1126; ^1^H-NMR (300 MHz, DMSO-d_6_); δ (ppm) 11.00 (s, 1H, NH), 10.35 (s, 1H, =CH-O), 7.24 (s, 1H, Ar-H), 6.48(s, 1H. furan). ^13^C-NMR (75 MHz DMSO-d_6_) δ (ppm) 150.0, 155.1, 143.4, 140.7, 138.1, 113.4, 116.2, 61.3; Anal. Calcd. for C_11_H_4_F_3_IN_2_O: C, 36.29; H, 1.11; N, 7.69; found: C, 36.07; H, 1.37; N, 7.74.

#### 4,5,6-trifluoro-2-(4-iodofuran-2-yl)-1-methyl-1H-benzo[d]imidazole (5n)

Yield: 81%; m.p.: 165-166 °C; FT- IR ν (cm^-1^): 3286, 3054, 2937, 1619, 1570, 1538, 1460, 1449, 1421, 1326, 1120. ^1^H-NMR (300 MHz, DMSO-d_6_); δ (ppm) 7.41 (d, 1H, =CH-O), 7.14 (s, 1H, Ar-H), 6.5 (s, 1H. furan), 3.75 (s, 3H, -CH_3_). ^13^C-NMR (75 MHz DMSO-d_6_) δ (ppm) 155.12, 150.10, 150.8, 147.49, 141.32, 138.6, 119.3, 109.3, 106.2, 100.25, 60.5, 30.4 Anal. Calcd. for C_12_H_6_F_3_IN_2_O: C, 38.12; H, 1.60; N, 7.41; found: C, 38.35; H, 1.74; N, 7.30.

#### 1-ethyl-4,5,6-trifluoro-2-(4-iodofuran-2-yl)-1H-benzo[d]imidazole (5o)

Yield: 73%; m.p.: 167-169 °C; FT- IR ν (cm^-1^): 3288, 3108, 2963, 1631, 1554, 1538, 1466, 1448, 1419, 1318, 1115; ^1^H-NMR (300 MHz, DMSO-d_6_); δ (ppm) 7.44 (s, 1H, =CH-O), 7.25 (s, 1H, Ar-H), 6.43(s, 1H, furan) 3.74 (m, 2H, -CH_2_), 1.57 (t, 3H, -CH_3_, J=18Hz).^13^C-NMR (75 MHz DMSO-d_6_) δ (ppm) 154.32, 150.3, 139.5, 141.22, 136.58, 125.1, 119.7, 106.2, 52.4, 36.6, 15.5; Anal. Calcd. for C_13_H_8_F_3_IN_2_O: C, 39.82; H, 2.06; N, 7.14; found: C, 40.03; H, 1.99; N, 7.23.

#### 4,5,6-trifluoro-2-(4-iodofuran-2-yl)-1-isopropyl-1H-benzo[d]imidazole (5p)

Yield: 78%; m.p.: 170-173 °C; FT- IR ν (cm^-1^): 3346, 3122, 3001, 1663, 1596, 1558, 1478, 1431, 1406, 1302, 1117; ^1^H-NMR (300 MHz, DMSO-d_6_); δ (ppm) 7.38 (s, 1H, =CH-O), 7.23 (s, 1H, Ar-H), 6.41(s, 1H. furan) 4.25(m, 2H, -CH_2_), 1.64(d, 3H, -CH_3_). 1.75(d, 3H, -CH_3_). ^13^C-NMR (75 MHz DMSO-d_6_) δ (ppm) 154.22, 149,77, 146.4, 141.12, 140.01, 119.23, 109.3, 106.1, 104.7, 45.3, 47.7, 21.4; Anal. Calcd. for C_13_H_8_F_3_IN_2_O: C, 41.27; H, 2.70; N, 6.87; found: C, 41.55; H, 2.52; N, 6.95.

## Biochemistry experiments

### Antioxidant activity

The *in-vitro* antioxidant free radical scavenging potential of the fuberidazole derivatives (**5a**-**5p**) was assessed utilizing a reduction method DPPH (2,2-diphenyl-1-picryl hydrazyl) with the slightly modified protocol(21, 22). The stock methanolic solution of DPPH was prepared (4 mg DPPH/4 mL MeOH) and further dilutions were also prepared at concentrations: 1.0, 2.0, 4.0, 8.0, 16.0, 32.0, 64.0, 128, 256, and 500 μg/mL. At 517 nm, the absorbance of sample solutions was determined. All sample solutions were produced and stored in the amber reagent bottle. As a positive control, *tert*-butyl-1-hydroxytoluene (BHT) was produced by dissolving 2 mg of BHT in methanol to obtain a mother solution (1000 µg/mL). The methanolic solutions of compounds **5a**-**5p** (2 mg each) were prepared to get a stock solution of 1000 µg/mL concentration. The serial dilution method was used to prepare test samples from this stock solution using methanol to attain the concentration ranges like DPPH. A 2.0 mL test compounds solution and 3.0 mL of DPPH solution (20µg/mL) were mixed, followed by vigorous shaking, and then kept undisturbed in a dark place for 30 minutes at ambient temperature. The absorbance at 517 nm has been determined employing methanol as a standard using a UV-Spectrophotometer. The fraction of the inhibited free radical DPPH was estimated utilizing the following equation:

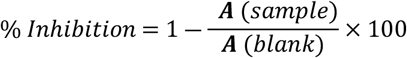

Where A blank refers to the control reaction absorbance.

## Cell culture

Human breast adenocarcinoma (MCF-7) was procured from ATCC (Manassas, VA, USA). These cells were preserved in RPMI-1640 (Roswell Park Memorial Institute) medium, which was accompanied with 10% FBS, streptomycin (10,000 mg/mL), and penicillin (10,000 U/mL) in 5% CO_2_ at 37 °C. Before further treatment, hypoxia cells were made by placing MCF-7 cells in a hypoxia incubator for 24 hours (1% O_2_ and 5% CO_2_)(23).

## WST cytotoxic assay

MCF-7 cells were exposed to a varied concentration of the vehicle (0.2% DMSO) or the synthesized compound (500 to 1 µM solution of DMSO) for control cells. After 48 hours of incubation with the examined drugs, the cell viability was determined using the WST-1 assay (Millipore)(24). In this assay, cellular mitochondrial dehydrogenase changed the tetrazolium salt WST-1 to formazan dye, whose concentration correlates with the viable cell in the culture. The absorbance at 440 nm was determined using a BioTek Synergy H1 plate reader (BioTek, Winooski, VT; USA). The percentage of viable cells was calculated using the following formula.

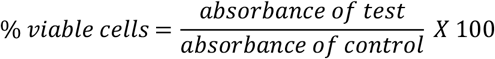

## DNA damage assay

The DNA-damaging effect of the substances was assessed using the EpiQuick in situ assay kit, which quantifies histone (H2AX) phosphorylation on serine 139. Seeding of A549/WM115 cells was carried out in 96-well plates at a concentration of 5000 cells / well and allowed to subsequent culturing under hypoxic conditions for 24 h prior to treatment. Subsequently, at the same culture conditions, the compounds were used to treat the cells/vehicles at concentrations ranging from 1C_50_ at the same culturing conditions. The DNA-damaging effect in the hypoxic cells was determined 4 hours after incubation, followed by fixation of cells and assay performance according to protocol. The DNA damage and intensity of color development in the samples were proportional. On a Synergy-H1 plate reader, the absorbance of the samples was determined at 430 nm. The damaged fraction of DNA was determined using the following expression:

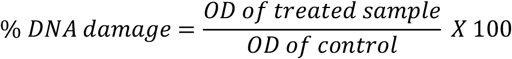

The optical density of samples is a measure of the absorbance of cells treated with test and control substances, as well as the optical density of the vehicle-treated control cells in the absence of test compounds.

## Apoptosis assay

This technique is based on using a proluminescent substrate containing DEVD to determine caspase-3/7 activity (Z-DEVD-aminoluciferin). Caspase Glo 3/7 assay [Promega] was employed based on the manufacturer’s instructions. Caspase cleavage liberates a resource for luciferase, which catalyzes the luciferase reaction and generates a bright signal. Caspase 3/7 activity was determined in hypoxic MCF-7 cells after 4, 24, and 48 h of treatment with the tested drugs. The luminescence at gain 135 was measured using a Synergy H1 and a Bio-Tek microplate reader(25).

## Computational methodology

### Properties of produced compounds based on chemo-informatics and ADMET

The cheminformatics and biological parameters of synthesized compounds (**5a**-**p**) were calculated by online servers, such as Molsoft and Molinspiration. Molinspiration was also used to observe the implication of Lipinski’s rule of five. Additionally, their ADMET properties were analyzed using the pkCSM online tool.

### Molecular docking

Docking simulations were performed on Autodock vina(26) following the reported literature(27-30) with slight modifications. The 3D structure of *human serum albumin* (PDB code: **1AO6**) was selected as target protein for the docking experiments with following parameters: size *x* = 20; size *y* = 20; size *z* = 20; center *x* = 25.2811; center *y* = 36.7115; center *z* = 23.1797.

### Nonlinear optical and optoelectronic studies

The Gaussian 09 software package, edition D.0131(31) was used to conduct all quantum chemistry calculations. Geometry improvements were performed at the B3LYP level(32) using the basis set 6-311+G(d,p)(33, 34). By demonstrating that the improved structures lacked fictitious frequencies (Nimg = 0), the harmonic-vibrational analytical frequency calculations verified that they are real minima. The energies shown are the zero-point free energies that have been modified. Stability analysis of the wave function indicated that the electronic ground states are closed-shell singlets, implying that the triplet energies are reliable. The Gaussian 09 program was also used to analyze the NLO response, the natural bond orbital (NBO), and the HOMO-LUMO pair. For NBO and NLO calculations, optimized geometries at the B3LYP/6-311+G(d,p) level were taken as a starting point. The electric molecular dipole moment (μ), first-order hyperpolarizability (β), and linear polarizability (α) were analyzed to evaluate the nonlinear response characteristics. The tensor components of polarizability (α) and hyperpolarizability (β) *(α*_*xx*_, *α*_*xy*_, *α*_*yy*_, *α*_*xz*_, *α*_*yz*_, *α*_*zz*_ and *β*_*xxx*_, *β*_*xxy*_, *β*_*xyy*_, *β*_*yyy*_, *β*_*xxz*_, *β*_*xyz*_, *β*_*yyz*_, *β*_*xzz*_, *β*_*yzz*_, *β*_*zzz*_) are obtained by the Gaussian output. The Gaussian output values (α and β) are transformed from atomic units (a.u) into electronic units (esu) (*α*; 1 a.u. = 0.1482×10^−24^ esu, and *β*; 1 a.u. = 8.6393×10^−33^ esu). When an isolated molecule is exposed to an incident electromagnetic wave’s applied electric field (E), microscopic polarizability may be produced in that molecule, the magnitude of which can be expressed as follows:

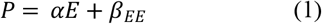

Where *P* and *E* represent tensor quantities, while α and β represent polarizability and hyperpolarizability, respectively.

Moreover, the average dipole moment (μ) can be defined as in equation (2)

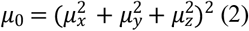

The polarizability (α) of a molecule can be defined as in equations (3)(35)

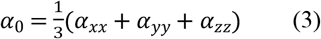

The anisotropy of polarizability is:

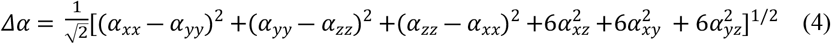

Equation (5) can be used to compute the components of first-order hyperpolarizability.

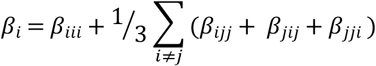

The first-order hyperpolarizability is a tensor component of the third rank, which can be explained by a 3 × 3 × 3 matrix(36). Then, the magnitude of the first hyperpolarizability can be determined using x, y, and z components as in equations (6) and (7).

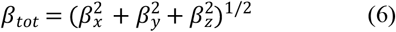

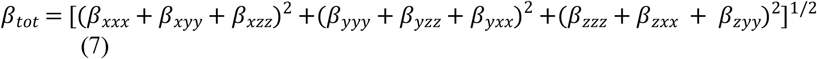

#### Statistical analysis of the data

For statistical analysis, ‘the student’s t-test was utilized. The significance level was set at the value of 0.05. The results are provided as mean standard deviation (± SD).

## Results and discussion

### Chemistry

To synthesize fuberidazole derivatives **5a**-**p**, a facile and green route was adopted in which 3,4,5-trifluorobenzene-1,2-diamine **1** was treated with furan-2-carbaldehyde derivatives **2** using PEG_400_ as a greener medium giving intermediate **3a**-**d** confirmed by elemental analysis and ^1^H and ^13^C NMR (see Table S1, Figs S1 and S8). The intermediate **3a**-**d** were then reacted with RCl **4** and sodium hydride in the presence of a minimum quantity of acetone as a solvent for five hours at room temperature, which afforded fuberidazole derivatives **5a**-**p** in good to excellent yields (Scheme 1). Some by-products were also obtained other than the desired product. Therefore, we purified them by preparative TLC and column chromatography. The structures of **5a**-**p** were assigned according to FT-IR, multinuclear (^1^H and ^13^C NMR) and CHN analysis. In the FTIR spectrum of **5a**-**p**, the absence of the -NH_2_ absorption band confirmed the formation of the imidazole ring. In the ^1^H NMR spectra of **5a**-**p**, the down-field singlets in the region 11.49 – 11.00 ppm were assigned to the NH ring proton. The peaks appeared in the range 3.75 – 1.55 are linked with -CH_2_ and -CH_3_ groups present in their respective compounds. All the aromatic protons of benzene moiety resonated between 7.45 – 6.25 ppm. A peak of one proton attached with carbon linked to the oxygen of furan moiety resonated downfield in the range of 7.87-7.46 ppm. The ^13^C NMR of **5a-p** showed maximum downfield shift for the carbon of furan attached with imidazole ring while azomethine carbon of imidazole moiety resonates at second-highest downfield shift in C-13 spectra. The other compounds were inveterate from their respective FTIR, multinuclear (^1^H, ^13^C) NMR spectra and CHN analysis.

**Scheme 1:**
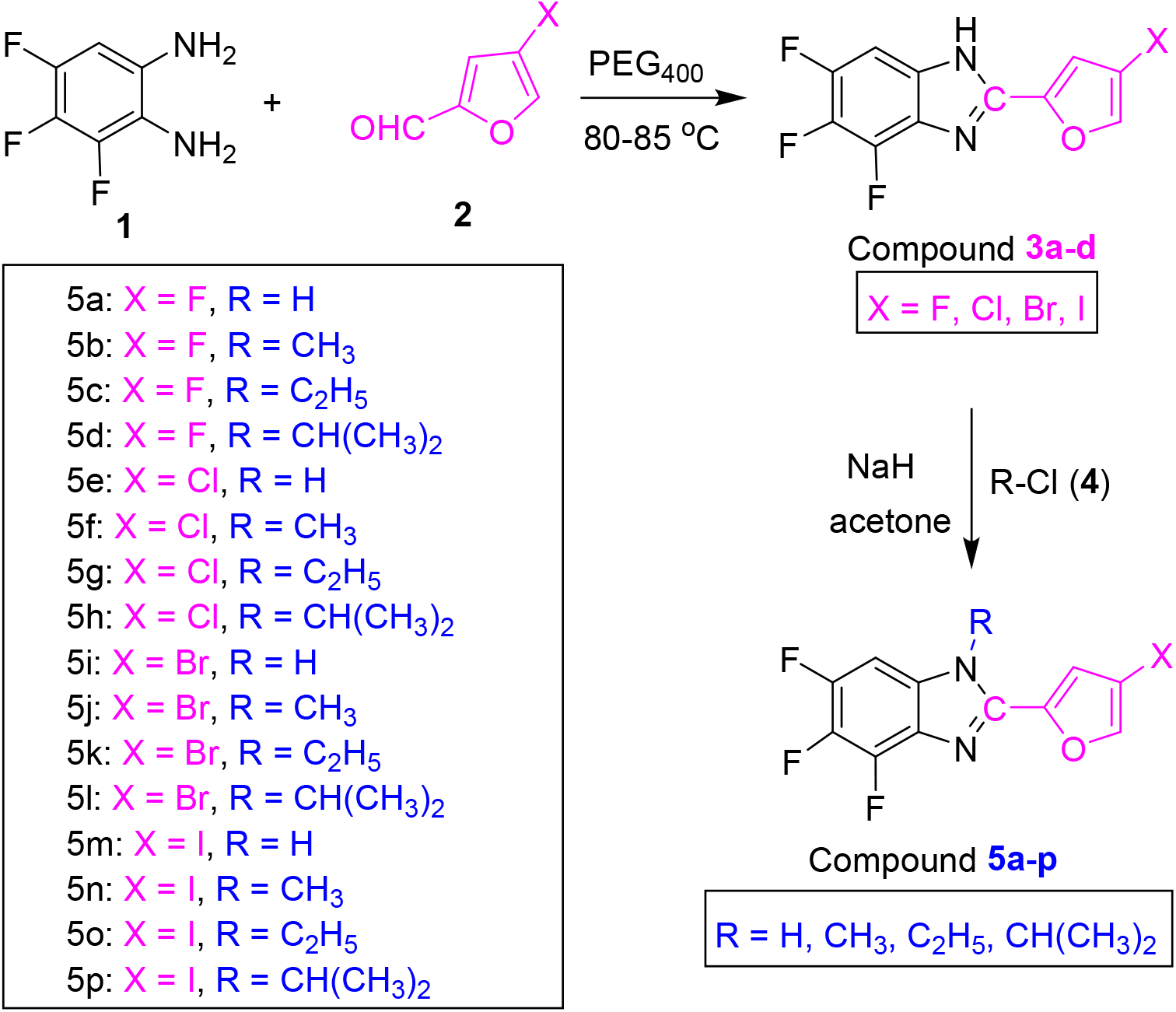
Synthetic scheme for synthesized compounds.

## Biological activities

### DPPH radical scavenging activity

The antioxidant potential was assessed using DPPH radical scavenging protocol having Butylated hydroxytoluene (BHT) as a positive control. The potential of the compoundc was determined by determining its electron-donating capacity toward DPPH, as shown by variations in the absorbance of a fluid of varying concentrations at 517 nm. The activity of DPPH radical scavenging of the fuberidazole derivatives’ linearly elevated with concentration. The findings on radical scavenging were described as the half-inhibition concentration (IC_50_), which is defined as the concentration which is necessary to scavenge 50% of DPPH radicals (Fig 1).

**Fig 1.**
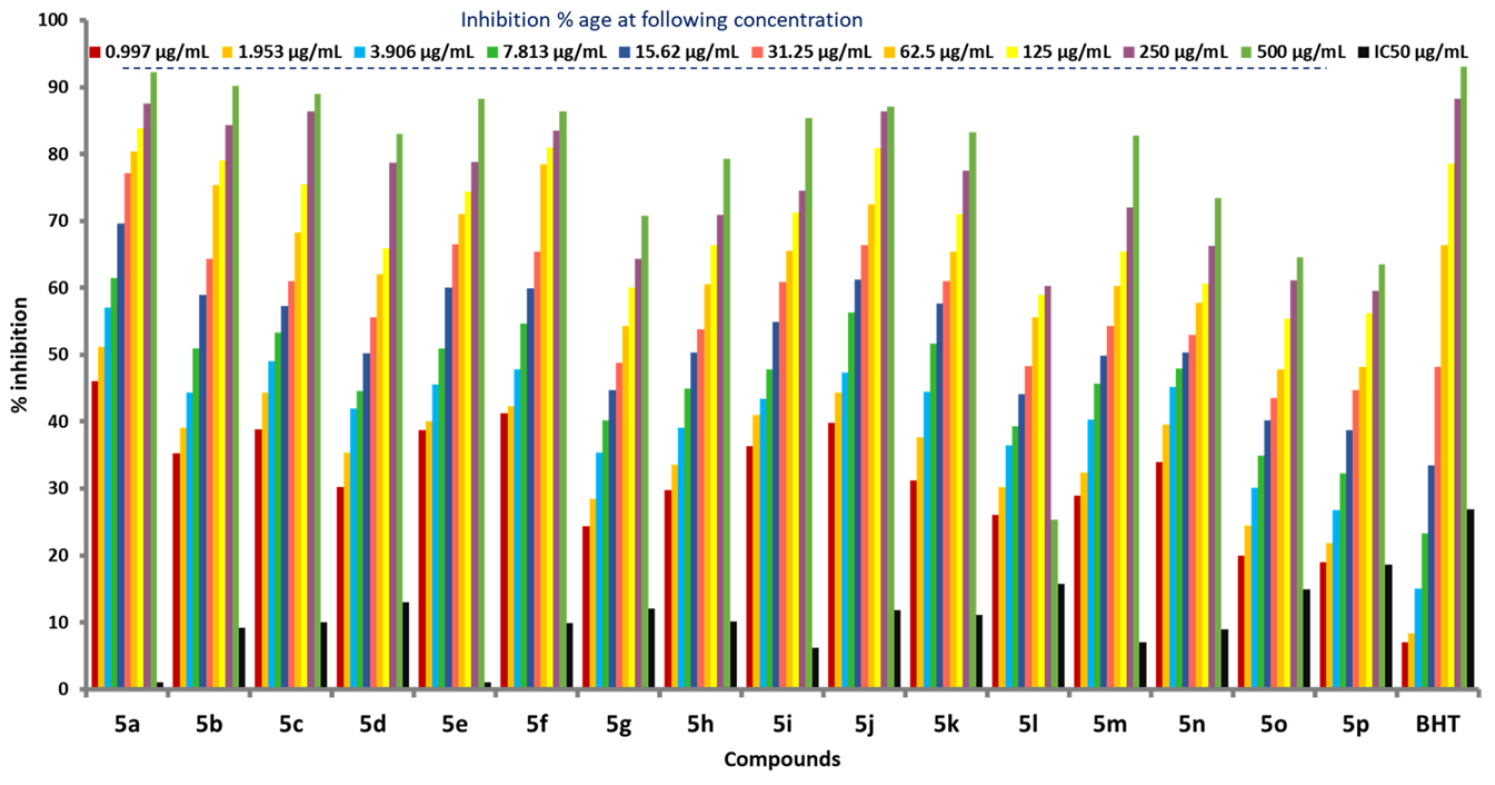
Antioxidant activity of compounds **5a-p**.

Fuberidazole derivatives **5a** and **5e** showed IC_50_ of 1.023 and 1.04 µg /mL, respectively, which seemed to be very promising and could be due to the presence of the halo group, which generally gives considerable biological activity. Concerning alkyl substituents, going to i-Pr from H-over CH_3_, and Et, quite a significant amount of compound is needed to show IC_50_ activity. Therefore, compounds substituted with H-atom (**5a, 5e, 5I, 5m**) showed better results than others. On the other hand, all fuberidazole derivatives (**5a**-**p)** exhibited mild to prominent antioxidant activity (18.67-1.023 µg /mL) because of the halogen substitution or the N atom, which is an important atom in many drug candidate molecules.

### Effect of compounds on cell proliferation of MCF-7 cells

We estimated the potential of synthesized fuberidazole derivatives on hypoxic MCF-7 cells. As revealed in Fig 2, compounds **5a, 5e, 5h**, and **5i** demonstrated the highest activity among the synthesized compounds (**5a**-**p**) in hypoxic conditions when compared to the hypoxic drug tirapazamine. The compound **5a** proved to be most active among **5a-p** against hypoxic cells, while the compound **5n** exhibited an inactive action. The exposure of hypoxic cells to **5a** at concentration ca. 100 µM resulted in an 80% reduction in cell survival. In contrast, compounds **5e** and **5h** showed cell survival of 53 and 56%, respectively, at the same concentrations. The **5n, 5o**, and **5p** have no obvious anticancer potential at any concentrations examined in hypoxic cells (Figs 2 and 3).

**Fig 2.**
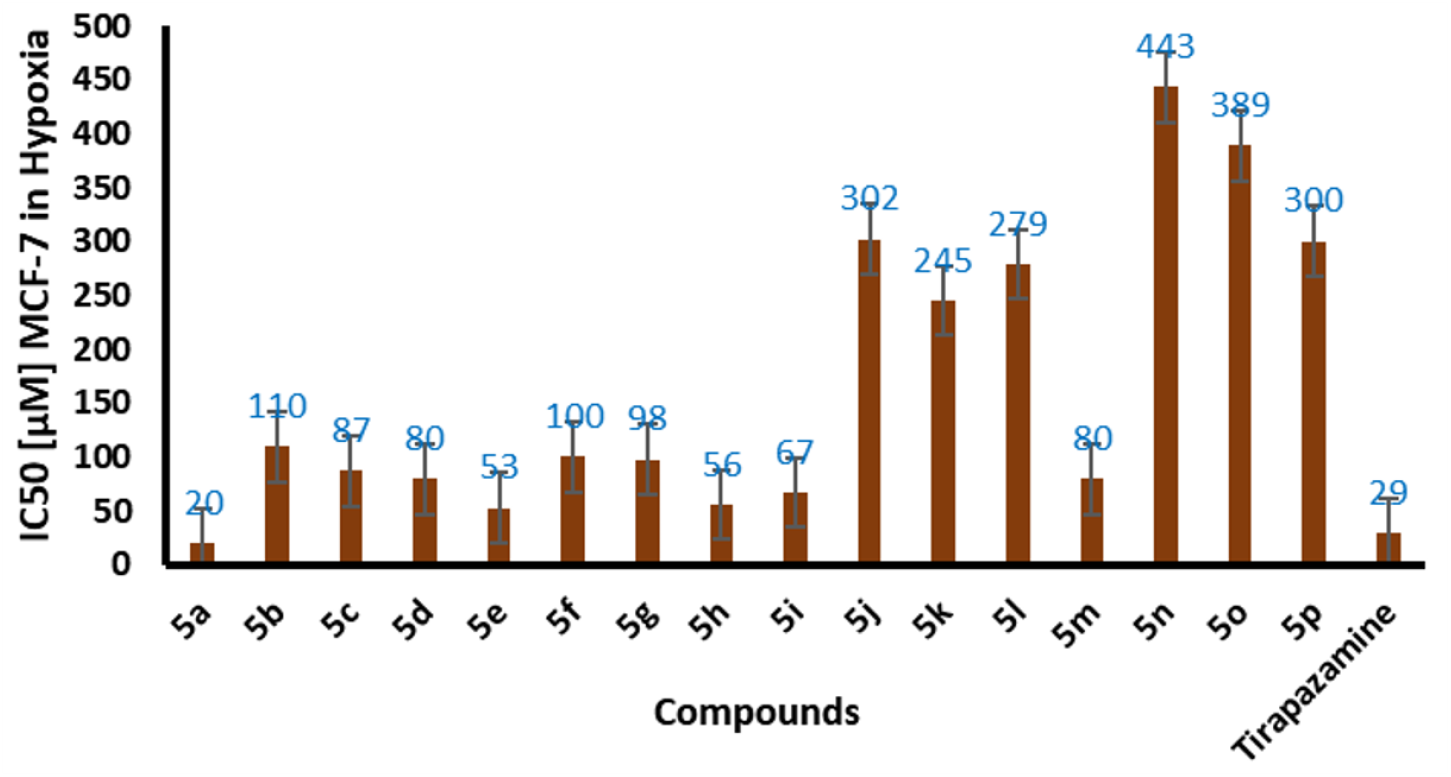
Fuberidazole derivatives 5a-p display a range in potency toward hypoxic MCF-7 cells. The WST-1 assay was utilized to examine the inhibitory impact of investigated drugs on cell growth during 48 h incubation period. The IC_50_ values (doses of examined substances that inhibit cell growth by 50% as compared to control cells) were determined and represented as the mean ± SD, n=3.

**Fig 3.**
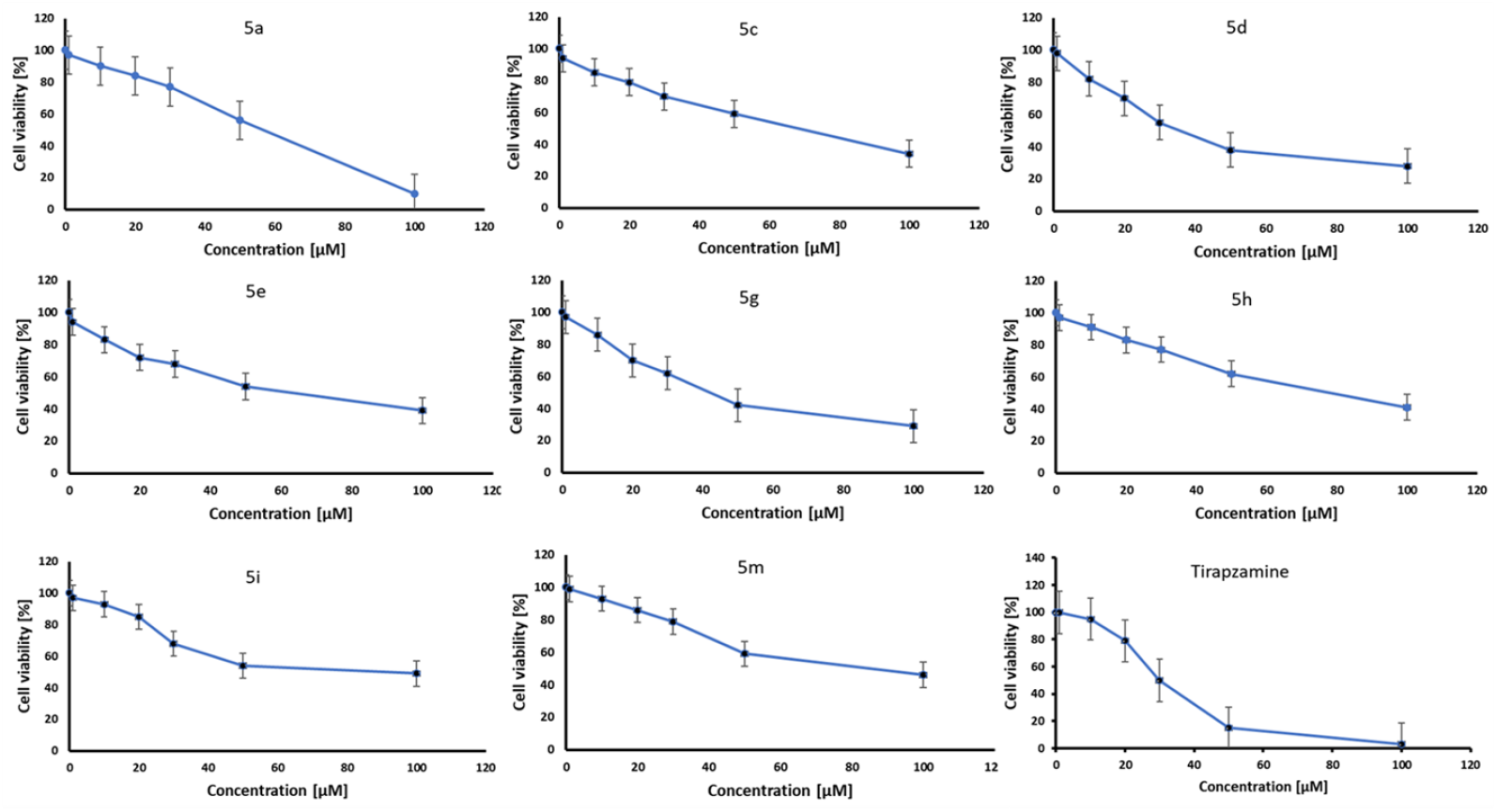
Fuberidazole derivatives affect MCF-7 cell viability in hypoxia. The WST-1 test was used to determine cell viability after 48 h of the treatment. The findings were given as a mean ± SD, n = 3.

Based on the results of the viability of hypoxic cells, the compounds (**5a**,**5c**-**e, 5g**-**i, 5m**) are highly active and selected for further testing.

### Potential of compounds 5a-p on DNA damage in hypoxic MCF-7 cells

In this regard, we calculated the phosphorylation of H2AX at Ser139 in hypoxic MCF-7 cells by exposing them for 4 h to the chosen compounds, tirapazamine and etoposide at the IC_50_ concentrations. The hypoxic cancer cells treatment with fuberidazole derivatives **5g, 5i**, and **5m** results in improved H2AX phosphorylation levels (Fig 4). Compound **5m** had the highest efficiency and enlarged the DNA damage by 1.5-fold in hypoxic cells compared to the control. In the hypoxic cells treatment with the fuberidazole derivatives **5d** and **5h**, the DNA damage was equivalent to that observed in control cells. Conversely, **5a** had no discernible effect on H2AX phosphorylation. Additionally, hypoxic cells exposed to tirapazamine caused DNA damage by 1.8-fold more than control cells. Because of these findings, we can conclude that compounds **5g, 5i**, and **5m** induced DNA damage in hypoxic MCF-7 cells at concentrations in the IC_50_ range.

**Fig 4.**
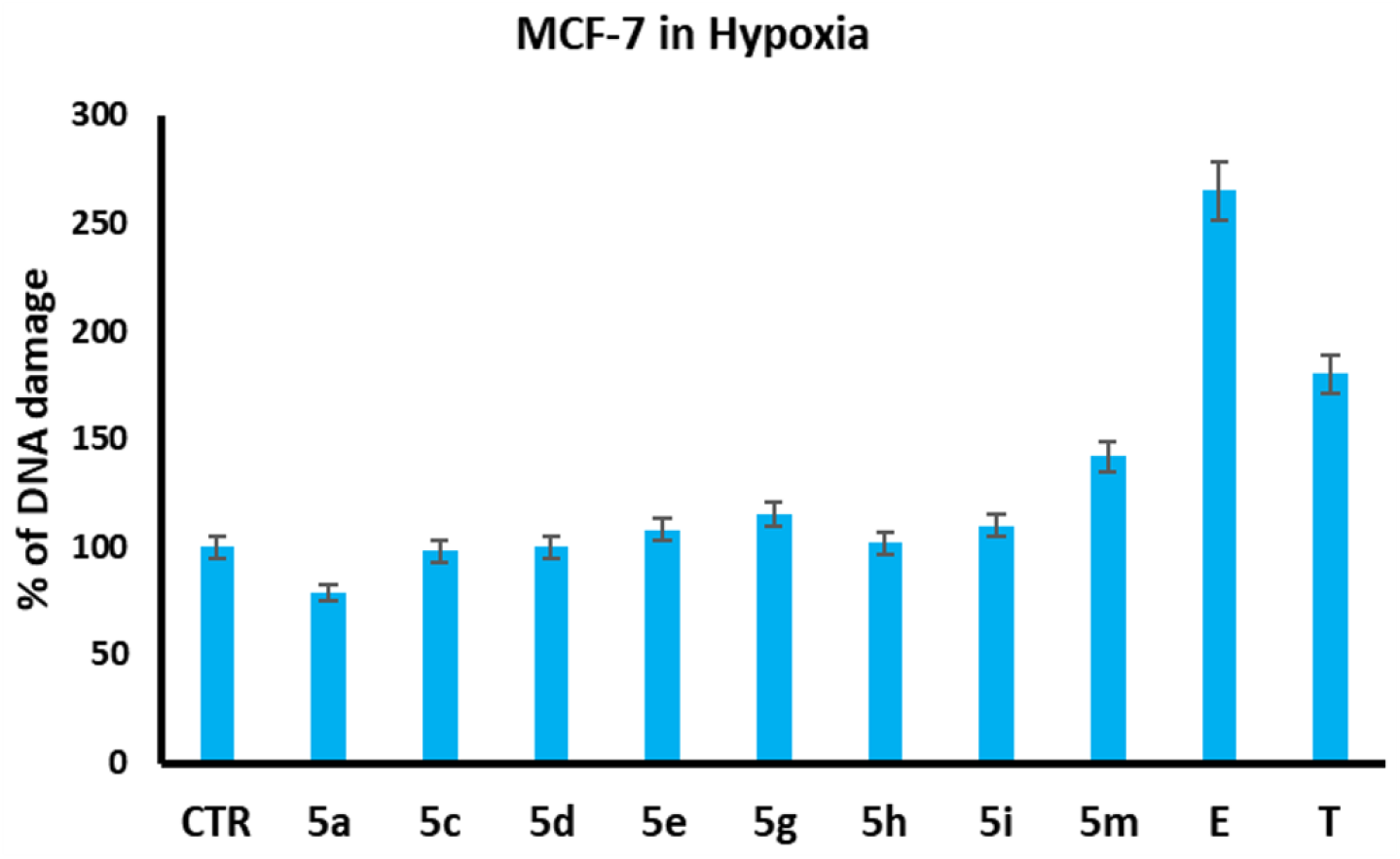
Effect of selected fuberidazole derivatives on DNA damage in hypoxic MCF-7 cells, CTR (control), E (etoposide), and T (tirapazamine). The control sample (CTR) included MCF-7 cells that had not been exposed to the substances under study. Statistical data: mean and standard deviation of trials are represented as n = 3.

### Potential of compounds on cell apoptosis

We selected compounds (**5a**,**5c-e, 5g-I**, and **5m**) for apoptotic activity using the IC_50_ values of fuberidazole derivatives. The activity of caspase 3/7 was investigated to assess the impact of apoptosis in the inhibition of hypoxic MCF-7 cell growth. Apoptosis assays were done in hypoxic MCF-7 cells treated with compounds (**5a, 5c-e, 5g-I**, and **5m**) at IC_50_ concentrations for different intervals. After 24 and 48 h, no elevation in the activity of caspase 3/7 was observed when hypoxic cells were treated with chemicals (**5a, 5c-e, 5g-I**, and **5m**) relative to control. In comparison, the treatment of hypoxic cells with tirapazamine for 24 h raised caspase 3/7 activity by seven times, while treating them for 48 h doubled its activity (Fig 5). The results indicated that the synthesized compounds (**5a, 5c-e, 5g-I**, and **5m**) inhibited MCF-7 cell growth rather than cell proliferation at the tested concentrations.

**Fig 5.**
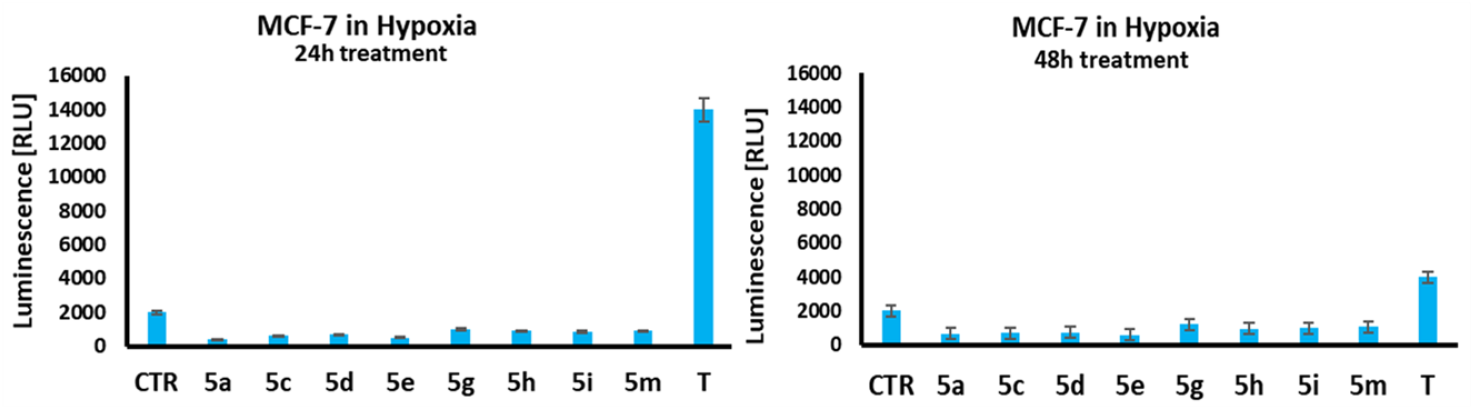
Exposure of selected fuberidazole derivatives to MCF-7 at hypoxia after 24 and 48 h. At the indicated concentrations, cells were cultured in the absence or presence (CTR, control) of the selected compounds and tirapazamine (T) as a control substance. Caspase 3/7 activity was determined after 24 and 48 h of therapy. The caspase’s activity was quantified by using a luminescence light unit [RLU] and represented as a mean ± SD, n = 3, * p < 0.05.

### A plausible structure-activity relationship of active fuberidazole derivatives

In the fuberidazole derivatives under investigation, the main pharmacophore group comprises the benzimidazole moiety as an essential receptor. The N–H group in imidazole may act as a critical site for the receptor. The presence of electron-withdrawing substituents to the adjacent furan ring’s *meta*-position could enhance the overall biological activity. In this regard, substituents F-, Cl-, Br-, and I-were used to investigating their effect on the pharmacological properties. The SAR is wisely explained in Fig 6.

**Fig 6.**
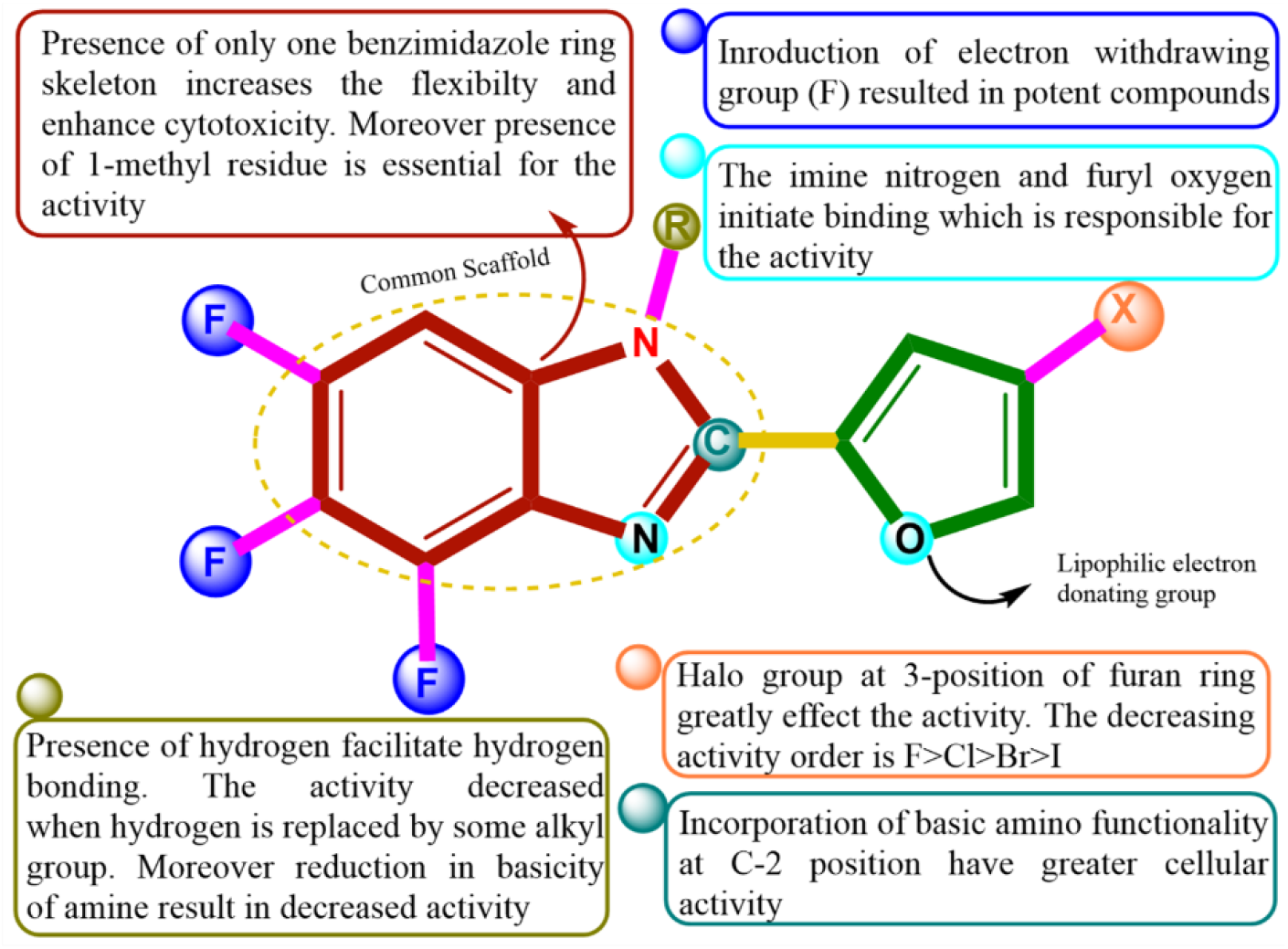
Display of structure-activity relationships in studied compounds.

### Chemo-informatics properties and Lipinski’s rule of compounds (5a-p)

The chemo-informatics characteristics of compounds (**5a**–**5p**), such as molecular weight, LogP, PSA, and molecular volume, were calculated and assessed by using computational methodologies listed in Table 1. The results confirmed that molecular weight (g/mol), molar volume (A^3^), and polar surface area (A^2^) of compounds **5a**-**5p** were correctly computed. All the calculated values for these substances were performed to be in agreement with the PSA (<89 A^2^)(37) standard value. Furthermore, the results of Lipinski’s rule of five (RO5) indicated that (**5a**–**5p**) compounds had appropriate HBD and HBA values, which substantially supported their drug-like activity. All compounds had molecular weights that were very similar to the standard limit (*<* 500 g/mol). The RO5 showed that molecules with low absorption have more an HBD value of more than five, MWT was greater than 500, a log*P* higher than 5, and an HBA value of more than 10.

**Table 1.**
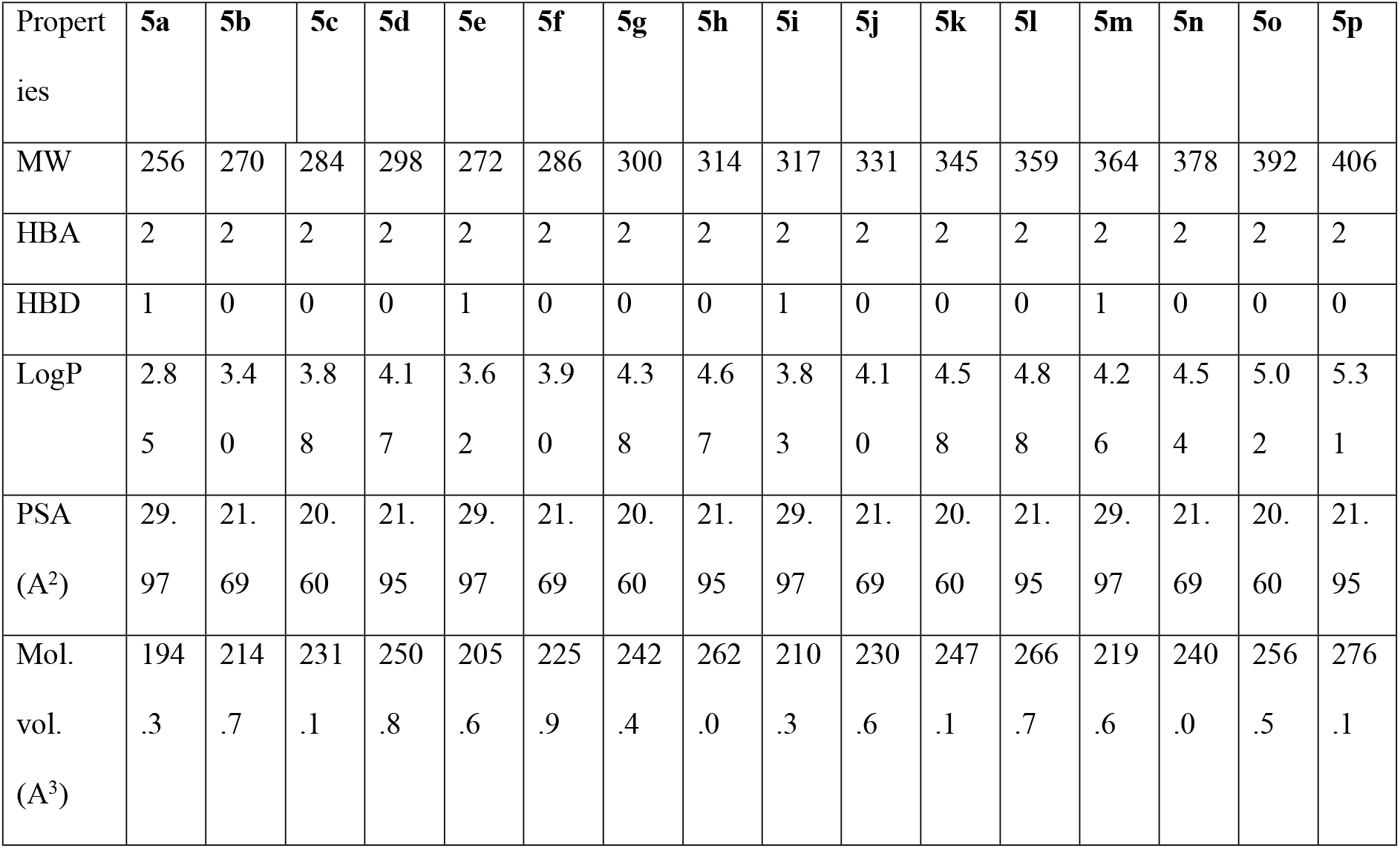
Chemo-informatics properties of fuberidazole derivatives.

### Pharmacokinetic properties of fuberidazole derivatives (5a-p)

The pharmacokinetics characteristics, such as Distribution, Metabolism, Absorption, Toxicity (ADMET), Excretion, and estimation, are thought to be the principal characteristics used to validate the efficiency of compounds. The ADMET properties of compounds have been calculated through the pkCSM online server (Table 2). It is stated that compounds with high absorption potential easily attack the target molecule by passively penetrating (crossing) the gut barrier(37). All compounds (**5a–p**) are highly water-soluble (WS) and exhibit a high intestinal absorbance (IA) when compared to the standard value (>30% abs). According to reports(38), a substance having an absorbance value of less than 30% is regarded to be poorly absorbed. Since all compounds have higher skin permeability (SP) values than the standard value (−2.5 log Kp) strongly recommends that they exhibit drug-like behavior. Moreover, the Central Nervous System (CNS) permeability and Blood-Brain Barrier (BBB) values of all compounds were within the normal range (>-2 to <-3 logPS and >0.3 to <-1 log BB), respectively. Only substances with a value larger than 0.3 log BB pass across the BBB, but those with less than -1 values are poorly dispersed to the brain. Likely, the substances with the LogPS values greater than -2 can penetrate the CNS, but those with a value less than -3 have difficulty spreading across the CNS.

**Table 2.** Pharmacokinetic assessment of fuberidazole derivatives (**5a-p**).

| ADMET Properties |  | 5a | 5b | 5c | 5d | 5e | 5f | 5g | 5h | 5i | 5j | 5k | 5l | 5m | 5n | 5o | 5p |
| --- | --- | --- | --- | --- | --- | --- | --- | --- | --- | --- | --- | --- | --- | --- | --- | --- | --- |
| **Absorption** | WS (log mol/L) | -2.927 | -2.957 | -2.961 | -3.048 | -2.94 | -3.043 | -3.064 | -3.103 | -2.948 | -3.04 | -3.06 | -3.10 | -2.95 | -3.04 | -3.06 | -3.104 |
|  | IA (% abs) | 85.6 | 90.8 | 90.5 | 88.1 | 84.4 | 89.75 | 89.48 | 87.58 | 84.38 | 89.68 | 89.41 | 87.52 | 85.02 | 90.32 | 90.05 | 88.15 |
|  | SP (log Kp) | -2.73 | -2.73 | -2.73 | -2.73 | -2.73 | -2.73 | -2.73 | -2.73 | -2.73 | -2.73 | -2.73 | -2.73 | -2.73 | -2.73 | -2.73 | -2.73 |
| **Distribution** | BBBP (Log BB) | 0.29 | 0.12 | 0.13 | 0.18 | 0.37 | 0.11 | 0.12 | 0.182 | 0.368 | 0.099 | 0.113 | 0.168 | 0.363 | 0.106 | 0.121 | 0.176 |
|  | CNSP (LogPS) | -1.99 | -1.82 | -1.85 | -1.72 | -1.85 | -1.76 | -1.80 | -1.68 | -1.85 | -1.76 | -1.79 | -1.68 | -1.85 | -1.76 | -1.80 | -1.68 |
| **Metabolism** | CYP3A4 | yes | no | yes | no | no | no | yes | yes | no | yes | yes | yes | no | yes | yes | yes |
|  | inhibitor |  |  |  |  |  |  |  |  |  |  |  |  |  |  |  |  |
| **Toxicity** | AMES toxicity | yes | no | no | no | yes | yes | yes | no | yes | yes | yes | no | yes | Yes | yes | no |
|  | ORAT (LD50) | 2.162 | 2.487 | 2.448 | 2.331 | 2.173 | 2.466 | 2.431 | 2.321 | 2.17 | 2.46 | 2.43 | 2.32 | 2.17 | 2.46 | 2.43 | 2.32 |
|  | Max. tolerat. dose | 0.313 | 0.042 | 0.098 | -<br>0.151 | 0.283 | -<br>0.009 | 0.059 | -0.18 | 0.28 | 0.009 | 0.05 | -<br>0.186 | 0.28 | -<br>0.007 | 0.061 | -0.18 |
|  | HT | yes | yes | yes | no | yes | yes | yes | no | yes | no | yes | no | yes | no | yes | no |
|  | SS | no | no | no | no | no | no | no | no | no | no | no | no | no | no | no | no |
| **Excretion** | TC(log ml /min/kg) | 0.63 | 0.45 | 0.56 | 0.49 | 0.63 | 0.45 | 0.56 | 0.49 | 0.66 | 0.47 | 0.62 | 0.51 | 0.65 | 0.46 | 0.61 | 0.50 |

“Abbreviation: WS: Water solubility, SP: skin permeability, IS: BBBP: blood-brain barrier permeability, intestinal solubility, ORAT: Oral Rat
Acute Toxicity, CNSP: CNS permeability, HT: Hepatotoxicity, SS: Skin Sensitization, TC: Total clearance.”

The compounds examined in this research displayed a high capability for crossing these barriers and binding to receptor molecules. The synthesized compound can inhibit the CYP3A4 enzyme (the isoform of cytochrome P450). The anticipated toxicity and excretion values indicated that these compounds exhibit the drugs-like behavior based on AMES toxicity, LD_50,_ maximum tolerated dose (MTD), and log ml/min/kg values. Additionally, the non-toxic and non-mutagenic behavior was evaluated using AMES toxicity prediction. Both negative and positive hepatotoxic behavior showed less sensitive and toxic effects. Most of the compounds displayed skin sensitizer and hepatotoxic properties. The ADMET properties results validated the prospective of these synthesized compounds to perform as lead compounds with minimal skin sensitivity and hepatotoxicity.

### Molecular docking

The interaction pattern was investigated using molecular docking, by which the optimal conformation of the fuberidazole derivatives in the target protein’s active site was determined, and the inhibitory efficacy against the target protein was assessed. Fig 7 illustrates the binding affinity score for inhibitors. All the inhibitors were docked in the proximity of the known active site (around the coordinates of TRP214). The AutoDock program was used to evaluate the optimum conformational position of the synthesized substances against *human serum albumin* (PDB code: **1AO6**). The bonding interaction pattern (hydrogen/hydrophobic) and minimum binding affinity values (kcal/mol) were used to evaluate the resultant docked complexes. Docking results discovered that **5e** exhibited the lowest value of binding affinity (−8.1 kcal/mol), while **5p** has predicted the binding affinity of -0.4 kcal/mol, which was found to be higher than the rest of the synthesized compounds. The supplemental data provide a list of all fuberidazole derivatives having docking complexes (Figs S41-S56).

**Fig 7.**
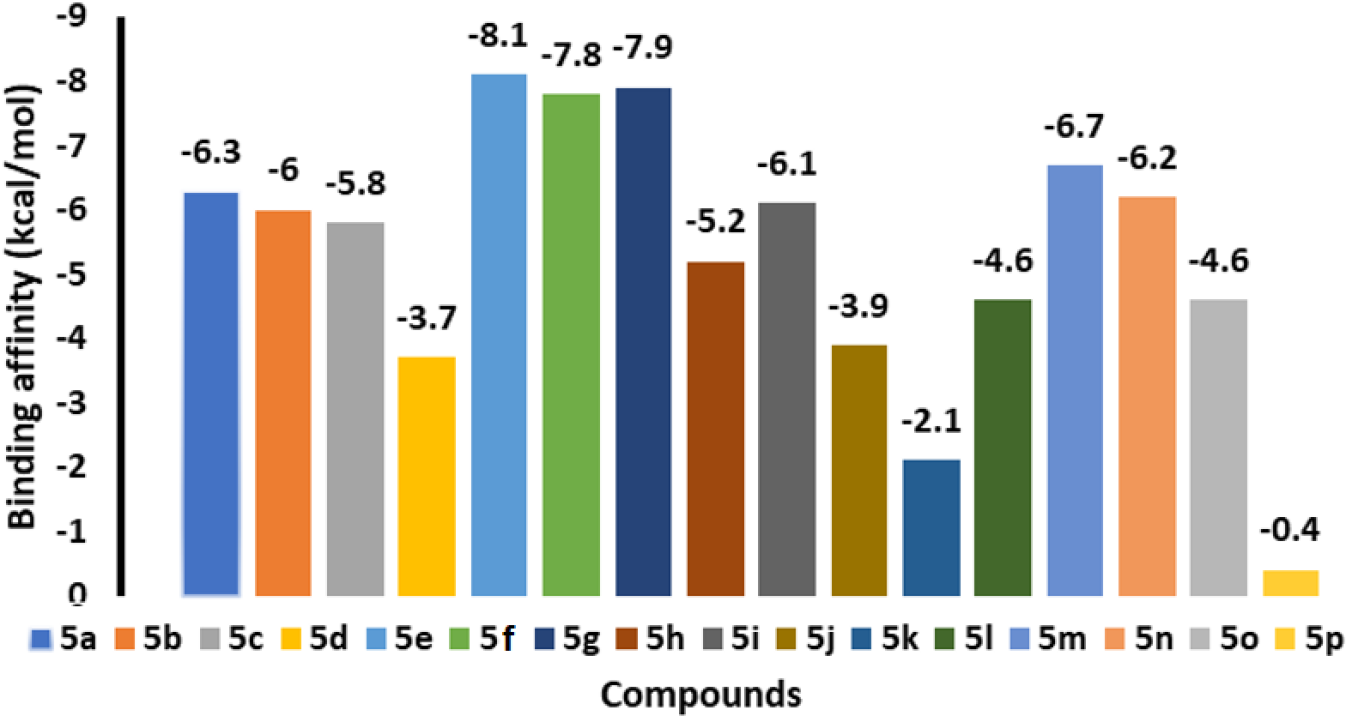
The graphical depiction of Binding affinity values of fuberidazole derivatives.

The docked complex was then analyzed further using hydrogen bonding, π--Alkyl, π-anion, π-Sulfur, π-sigma, and Van der Waals interactions. Docking analysis demonstrated that all docked substances interacted with the active binding area of *human serum albumin* [30]. On the other hand, compound **5e** had the lowest binding affinity (−8.1 kcal/mol) and the best fit into the active pocket of *human serum albumin*. As seen in the 3D and 2D images of compounds **5e** and **5g** (Figs 8-9), hydrogen bonding and hydrophobic interactions dominate the interaction between the formed substances and the target protein (Figs 10-11). ARG 222 from the active pocket of the protein has been observed to form hydrogen bonds. It is noted from the data that the residues ILE290, LEU 219, TYR150, HIS242, LYS199, LEU238, and ALA291 have been involved in hydrophobic interactions in the protein-ligand complex. Compound **5e** also exhibited Van-der-Waal interactions with the *human serum albumin*.

**Fig 8.**
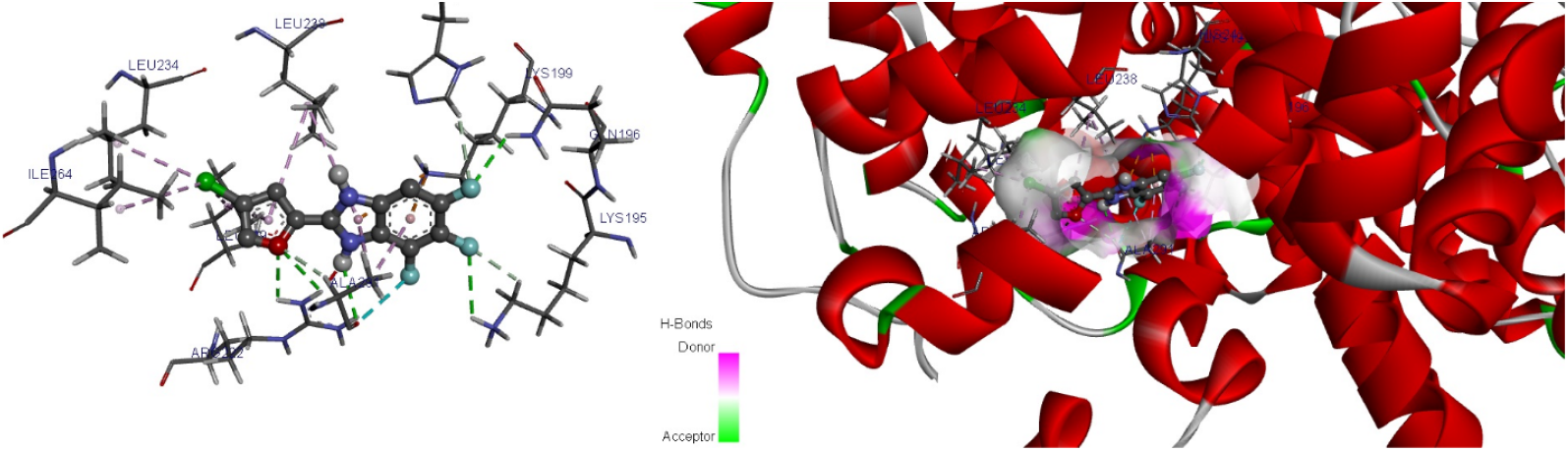
3D depictions of **5e** docking in the active site of *human serum albumin*.

**Fig 9.**
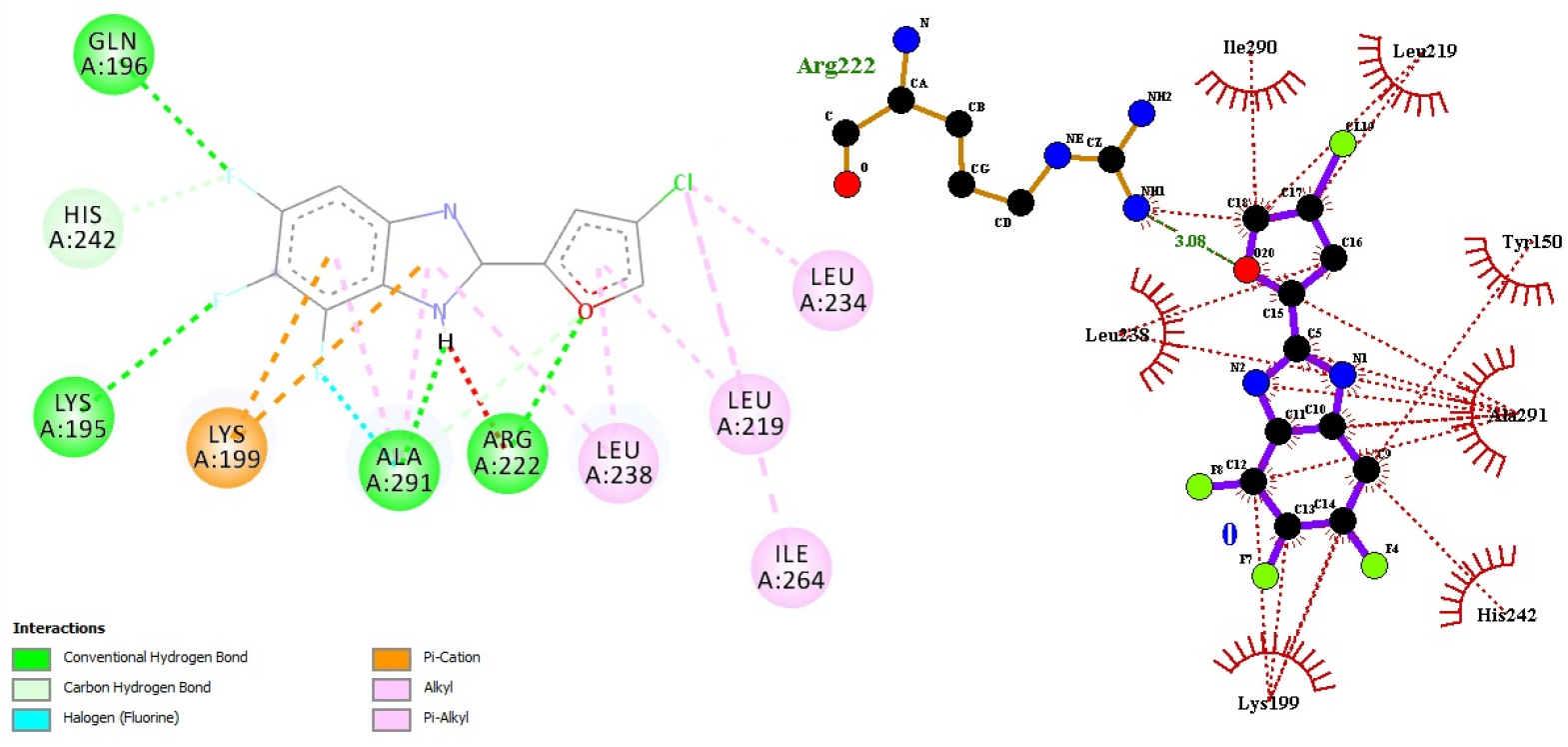
2D depiction of docking of **5e** in the active site of *human serum albumin*.

**Fig 10.**
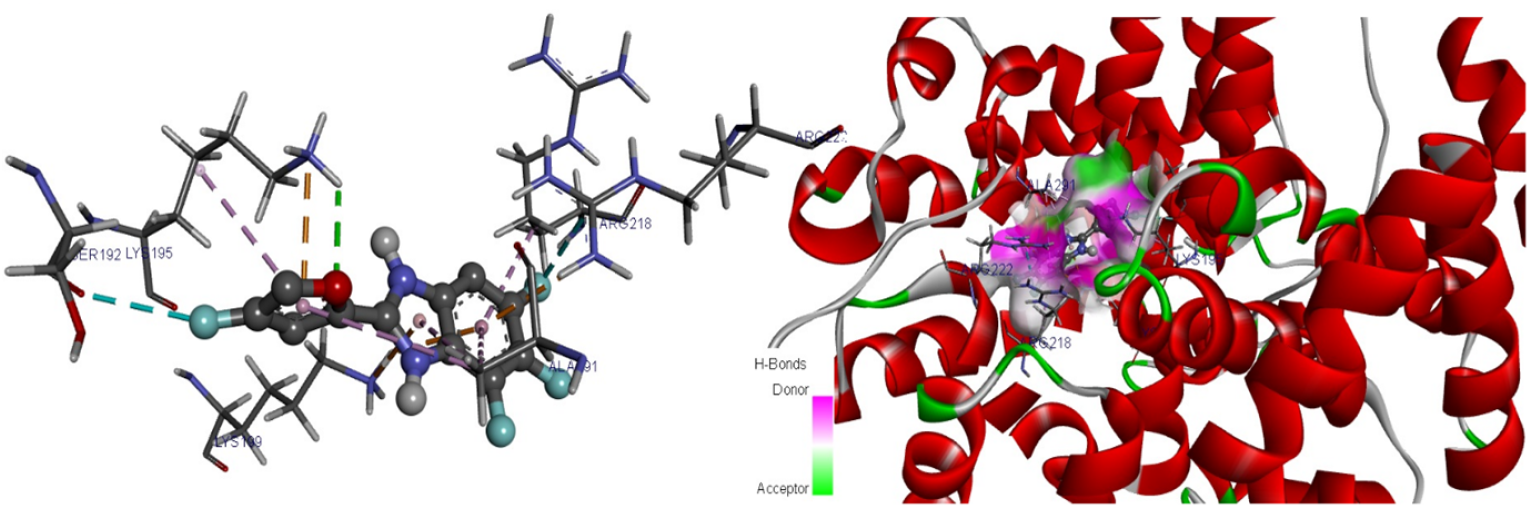
3D depictions of docking of **5g** in the active site of *human serum albumin*.

**Fig 11.**
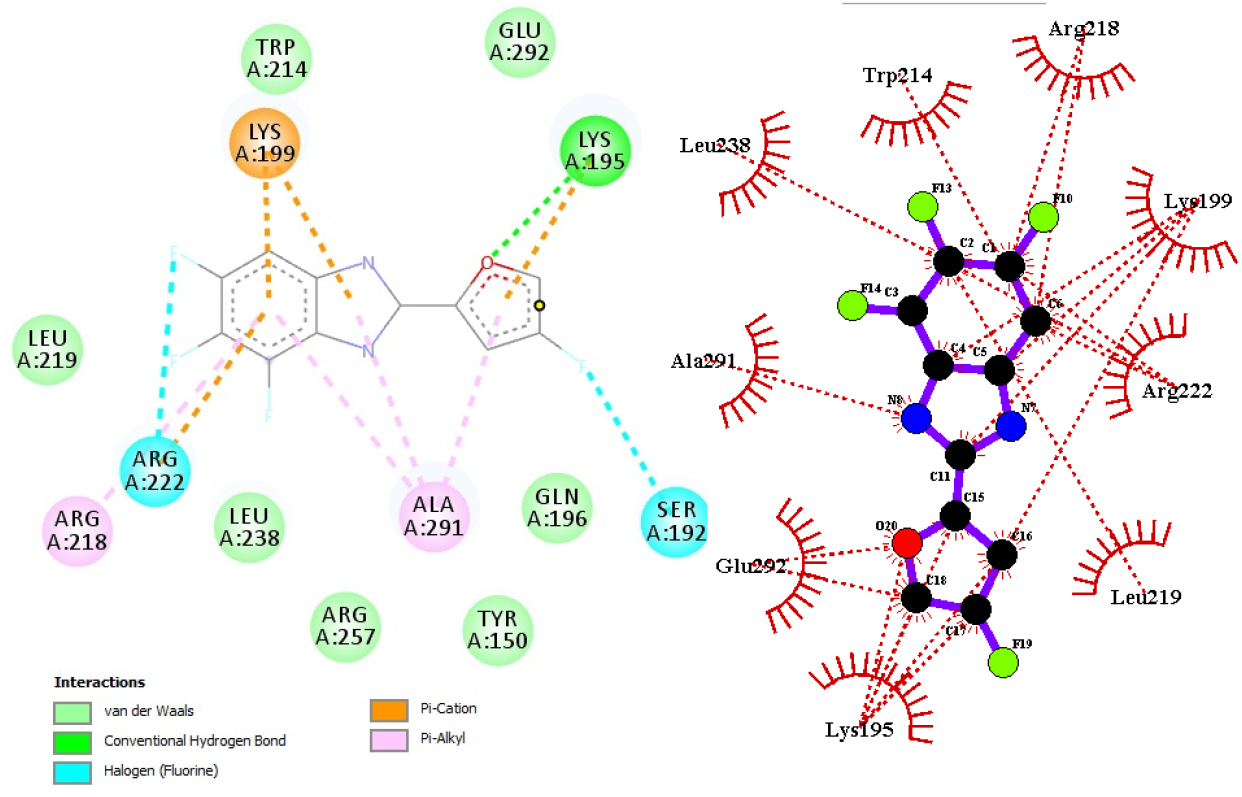
2D depiction of docking of **5g** in the active site of *human serum albumin*.

Docking results revealed that the compound **5g** was tightly bonded to the active site of the *human serum albumin* (binding affinity -7.9 kcal/mol) and established three ordinary hydrogen bonds with PRO C:303, PRO C:306, and GLU C:314. Van-der-Waal interactions were observed by **5g** with several amino acid residues in the binding pocket of *human serum albumin*. Overall, the docking study determines the reported compounds’ high affinity for *human serum albumin* (PDB code: **1AO6**), indicating that they might be regarded as potent suppressors in the future of drug discovery.

## DFT analysis

At the (U)B3LYP/6-311+G(d,p) level, the optoelectronic and first-order NLO response properties of a series of fuberidazole derivatives **5a**-**p** (Scheme 1) with different substituents at the N-atom of imidazole ring (−H, -Me, -Et, and -i-Pr) and the 3-position of furan ring (−F, -Cl, and –Br), respectively. We first discuss triplet energies and HOMO-LUMO gaps. Secondly, dipole moments, polarizabilities, and first-order hyperpolarizabilities are discussed. Finally, NBO analysis is carried out to investigate the donor-acceptor interaction.

### Triplet energies

Quantum chemical calculations of the gas-phase triplet energies revealed that all fuberidazole derivatives (**5a**-**d**) have lower triplet energy than their bromo- and chloro-substituted ones (Fig 12). This could be interpreted as fluorination having the effect of lowering the triplet energy. Bromo-substituted compounds (**5i**-**l**) display the highest triplet energies. In the case of alkyl substituents on the N-atom of the imidazole unit, methyl-substituted compounds (**5b, 5f, 5j**, and **5n**) showed the lowest triplet energies within each series. The triplet energies have been increased when going from methyl to isopropyl over ethyl substituent, which could be due to the steric repulsion. The increase in the steric bulk at the N-atom of the imidazole unit reduces the planarity of the compound, which weakens π-conjugation and increases triplet energy. Interestingly, compound **5b** showed the lowest triplet energy (2.29 eV). However, when the imidazole moiety’s benzene ring and furan ring are unsubstituted, the triplet energy is 2.38 eV, which is significantly higher than that of substituted compounds. Introducing electron-donating substituents (alkyl groups) on the N-atom of the imidazole (acceptor site), at the same time, introducing electron-withdrawing substituent on the furan ring (donor side) leads to a significant decrease in triplet energies. In summary, with proper choice of the electron-donating substituent on the acceptor unit and electron-withdrawing substituent on the donor unit, it could be possible to control the triplet energies of this series of compounds, possibly due to the push-pull effect.

**Fig 12.**
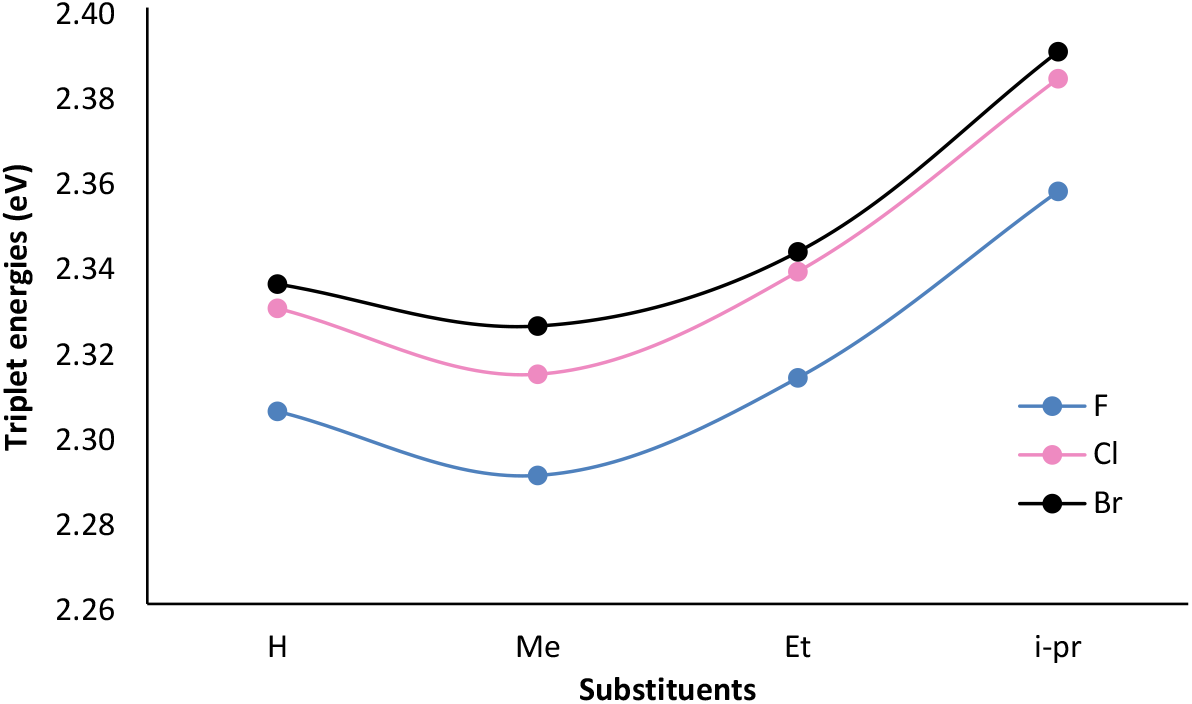
Triplet energies (eV) at the (U)B3LYP/6-311+G(d,p) level.

### HOMO-LUMO energy gaps

The electronic, optical, and spectroscopic properties of fuberidazole derivatives can be analyzed with the help of frontier molecular orbitals (FMOs). FMOs are also a form of electrical conductivity and can provide information about intramolecular charge transfer. The lower the HOMO-LUMO energy gap, the greater the probability of charge transfer. The results of the orbital analysis showed that HOMO-LUMO gaps for all compounds range from 4.88 – 4.33 eV (Fig 13). The smallest ΔE_H-L_ is found for compound **5a** (4.33 eV). However, the energies of HOMO and LUMO are not significantly affected by changing the substituents. The excitation is mainly ππ*.

**Fig 13.**
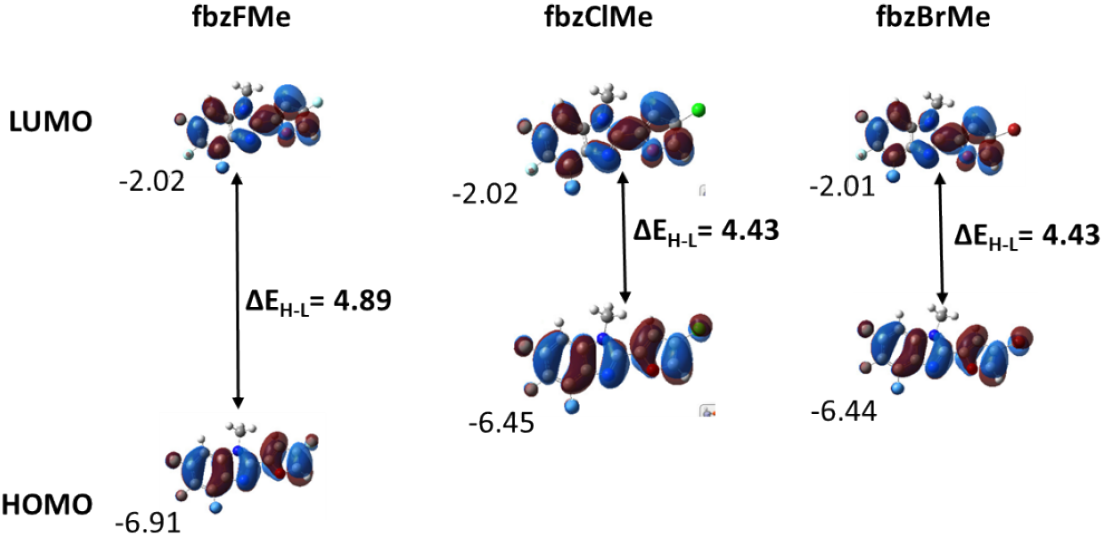
The molecular orbitals and respective energies (eV) of methyl-substituted fuberidazole compounds at the B3LYP/6-311+G(d,p) level. LUMO and HOMO denote the lowest and highest occupied molecular orbitals, respectively.

### Dipole moments

The molecular dipole moments are used to indicate the charge transport along the molecule(39). The gas-phase calculated dipole moments range from 5.54 – 6.49 Debye (Fig 14). Interestingly, dipole moments increased when the electronegativity of the halogen atom is decrease at the furan unit. In addition, a significant increase in dipole moments is observed by changing the substituent on the N-atom. The highest dipole moment is observed for **5l** and the lowest for **5a**.

**Fig 14.**
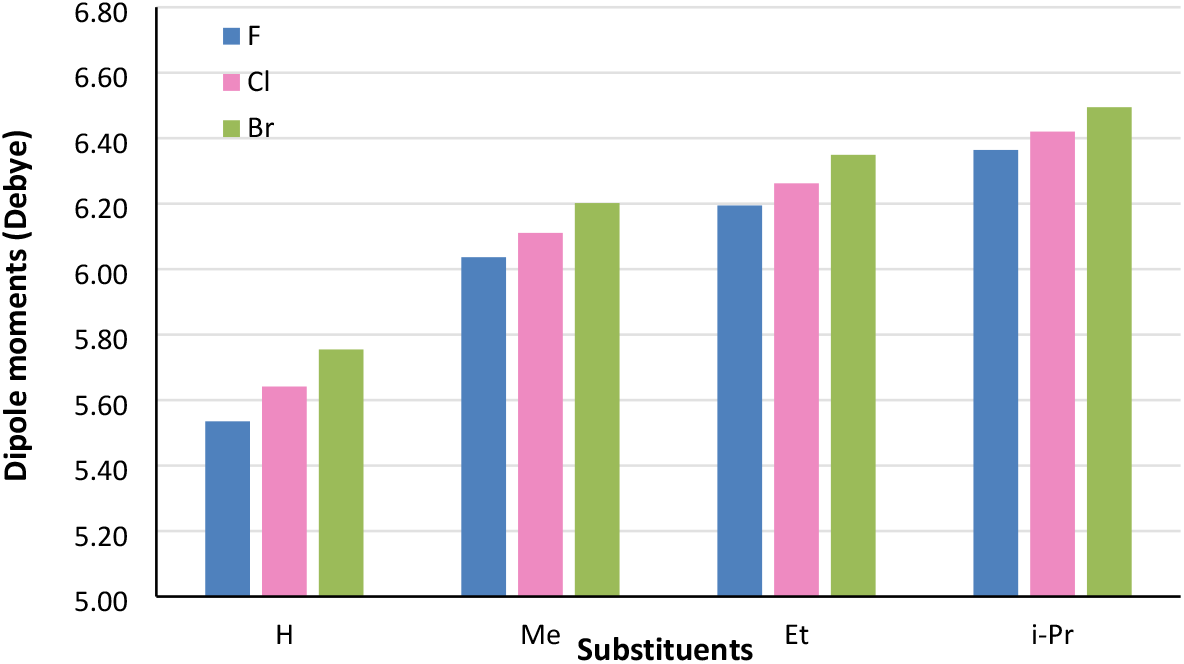
Dipole moments (Debye) in the gas phase at B3LYP/6-311+G(d,p) level for fuberidazole compounds (**5a-p**).

### Polarizability and first-order hyperpolarizabilities

The polarizability refers to the magnitude of electron density distortion and responsiveness of a system when a static electric field is applied externally. It was reported that particularly large values of hyperpolarizability are a key requirement for a molecule to be a good NLO material. The polarizability (α_ave_) and first-order hyperpolarizability (β_tot_) showed a similar trend to what was observed for dipole moment. Both above properties decrease with increasing the electronegativity of the halogen atom and increase by the introduction of the bulky electron-donating alkyl substituent at the N-atom(40). This can be explained as electron-donating groups increase the induced ring current, which can then be responsible for the enhanced NLO response. However, the effect of substituents on α_ave_ is less pronounced. The α_ave_ range from 2.37 – 3.18 × 10^−23^ esu and β_tot_ range from 49.11 - 69.89 × 10^−31^ esu. The highest value of α_ave_ and β_tot_ is observed for **5l**. The results support the hypothesis that polarizability and first-order hyperpolarizabilities can be tuned by changing the substituents on the donor and the acceptor units (Fig 15). Moreover, these fuberidazole derivatives could potentially be good candidates for NLO applications.

**Fig 15.**
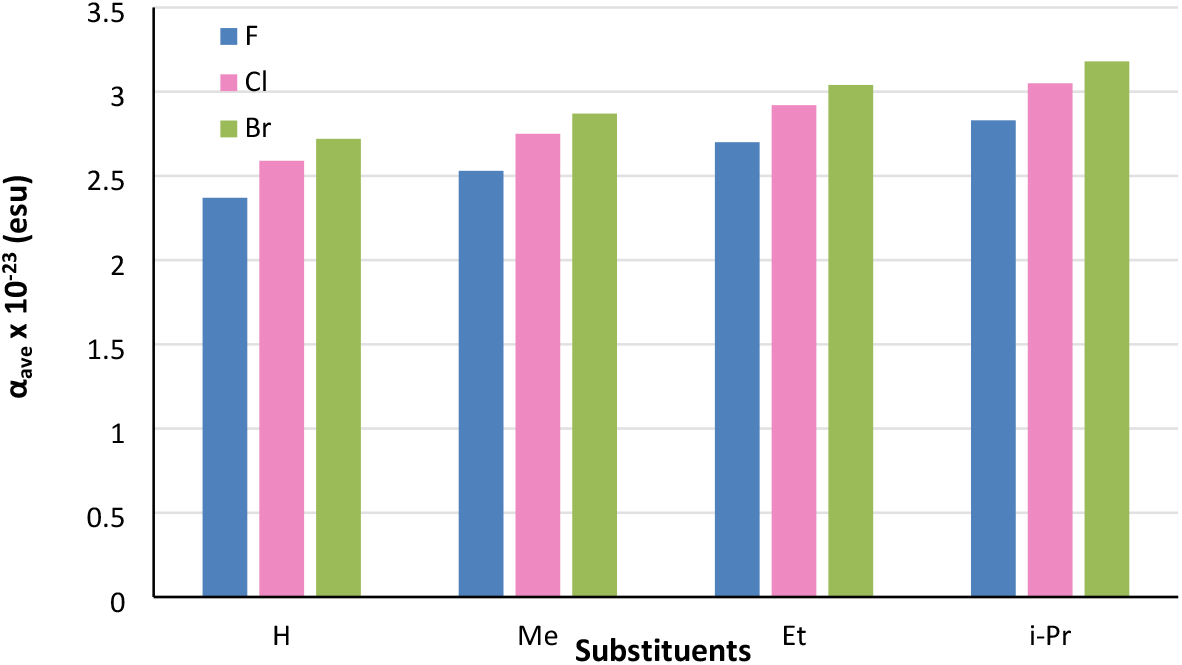
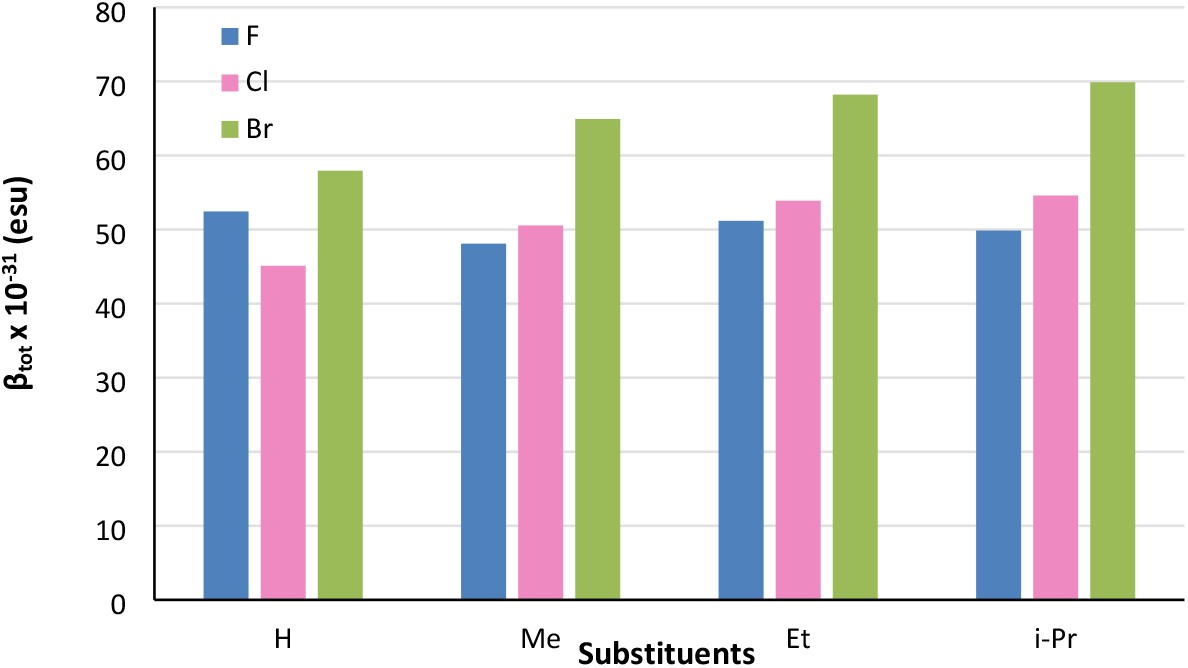
The α_ave_ × 10^−23^ (esu) and β_tot_ × 10^−31^ (esu) at the B3LYP/6-311+G(d,p) level for fuberidazole derivatives (**5a-p**).

### Natural bond orbital (NBO) analysis

The natural bond orbital (NBO) method of Weinhold provides information about the interaction within the different parts of the molecules, namely donor and acceptor units(41). The charges on different sites of the molecule were calculated by NBO to analyze the donor-acceptor interaction in the molecule. In fuberidazole, the benzimidazole unit, when substituted with electron-donating groups, is known to act as an acceptor moiety in push-pull chromophores(42, 43). Our results show similar results as the benzimidazole unit is acting as an acceptor, and the furan unit is acting as a donor in fuberidazole (Fig 16). The strong donor-acceptor push-pull interaction is observed in **5l** among the series of compounds, which might be responsible for increased α_ave_ and β_tot_ values.

**Fig 16.**
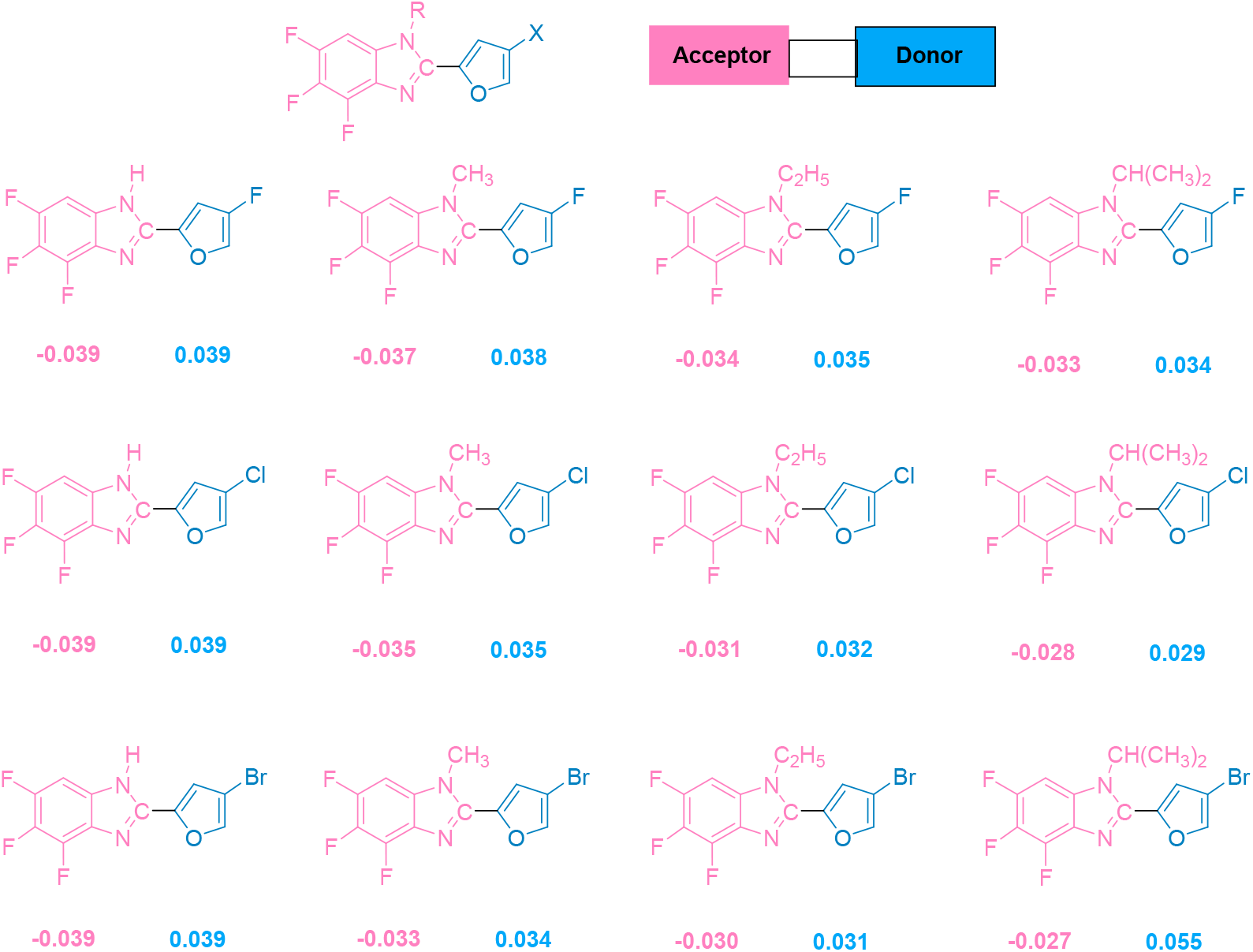
NBO charges calculated at the B3LYP/6-311+G(d,p) level illustrating donor-acceptor interaction in fuberidazole derivatives **5a-p**.

### A thought to extend conjugation by π-bridge

As it is known that the NLO response of materials can be enhanced by placing a π-bridge in between the donor and acceptor units(44). We designed three compounds with different π-bridges and calculated α_ave_ and β_tot_ for them at the B3LYP/6-311+G(d,p) level. It is found that when the C=N bridge is placed, a significant enhancement in β_tot_ is observed. On the other hand, when benzene is inserted as a π-bridge, no improvement in β_tot_ is observed. This reflected that the benzimidazole unit is separated by a π-bridge with a furan unit, which could display significantly enhanced NLO response characteristics. However, these compounds are only designed computationally (Fig 17). In the future, we have a plan to synthesize and investigate the practical applications of synthesized compounds.

**Fig 17.**
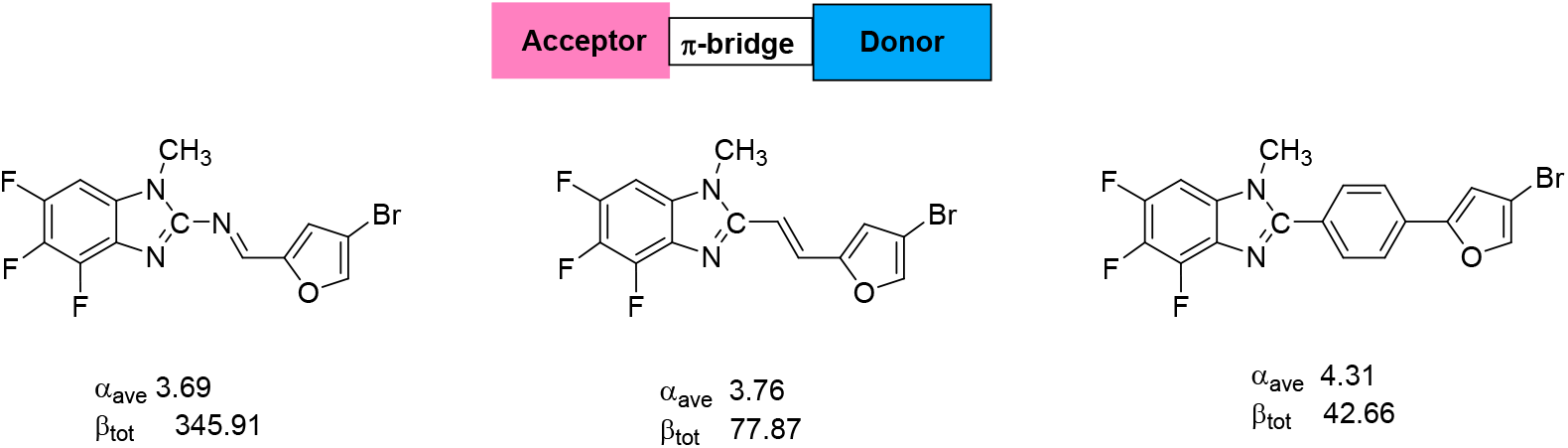
α_ave_ and β_tot_ of fuberidazole derivatives with π-bridge in between donor and acceptor units. The values for α are given in 10^−23^ esu and for β_tot_ in 10^−31^ esu.

## Conclusion

The biochemical research conducted in this study enabled us to establish certain conclusions regarding the biological impacts of fuberidazoles. We have described the synthesis, in-vitro, and in silico screening of sixteen fuberidazole derivatives as potential new anticancer candidates. All compounds showed mild to prominent antioxidant activity, but few (**5a, 5e, 5I**, and **5m**) showed better results than others. The in-vitro anticancer potential, apoptosis, proliferation, and DNA destruction experiments on selected hypoxic cancer cells were also described. Out of sixteen, eight compounds (**5a**,**c**,**d**,**e**,**g**,**h**,**i**,**m**) displayed good cytotoxic properties. This moderate relation between antioxidant and anticancer activities of tested substances implies that anticancer activity in the examined MCF-7 cells may be attributed to their antioxidant properties. SAR, pharmacochemical strength and the mode of interactions responsible for the activity have also been examined via in silico studies (against *human serum albumin*) using chem-informatics. Furthermore, molecular docking research was also conducted to calculate binding energy and protein interaction. Our docking studies established the described compounds’ high affinity for *human serum albumin* (PDB code: **1AO6**). As a result, these molecules may be regarded powerful inhibitors throughout the drug development process in the future. Finally, our results of quantum chemical calculations show that most electronegative F atom and least bulky alkyl groups at the acceptor and the donor sides, respectively, can decrease the triplet energies. It is found that the least electronegative atom and the bulkiest electron-donating substituent on the donor and the acceptor sides, respectively, show relatively highest values for β_tot_ (69.89 × 10^−31^ esu), α_ave_ (3.18 × 10^−23^ esu), and dipole moment (6.49 Debye). The compound **5l** showed the best results amongst the series of compounds. The donor-acceptor sites were also identified by the NBO analysis. Therefore, depending on the choice of application, one can tune the optoelectronic properties by properly placing substituents on the donor and the acceptor units. Noteworthy, the nonlinear response characteristics of compounds (**5a-p**) are investigated only; theoretically, one needs to confirm these results experimentally because these compounds could potentially be used as future organic linear and NLO materials.

## Acknowledgments

We are highly acknowledged for and Islamia University Bahawalpur for the synthesis and characterization of fuberidazole derivatives, Uppsala Multidisciplinary Center for Advanced Computational Science (UPPMAX) and Swedish National Infrastructure for Computing (SNIC) at the National Supercomputer Center (NSC), Linköping for computational resources and University of Jeddah Saudi Arabia for the bioactivities of fuberidazole derivatives.

